# The logic of recurrent circuits in the primary visual cortex

**DOI:** 10.1101/2022.09.20.508739

**Authors:** Ian Antón Oldenburg, William D. Hendricks, Gregory Handy, Kiarash Shamardani, Hayley A. Bounds, Brent Doiron, Hillel Adesnik

## Abstract

Recurrent cortical activity sculpts visual perception by refining, amplifying, or suppressing incoming visual signals. Despite the importance of recurrent circuits for cortical processing, the basic rules that govern how nearby cortical neurons influence each other remains enigmatic. We used two-photon holographic optogenetics to activate ensembles of neurons in Layer 2/3 of the primary visual cortex (V1) in the absence of external stimuli to isolate the impact of local recurrence from external inputs. We find that the spatial arrangement and the stimulus feature preference of both the stimulated and the target ensemble jointly determine the net effect of recurrent activity, defining the cortical activity patterns that drive competition versus facilitation in L2/3 circuits. Computational modeling suggests that a combination of highly local recurrent excitatory connectivity and selective convergence onto inhibitory neurons give rise to these principles of recurrent activity. Our data and modeling reveal that recurrent activity can have varied impact, but a logic emerges through an understanding of the precise spatial distribution and feature preference of the multicellular pattern of activity.

## Introduction

Visual perception involves the coordinated activity of thousands of neurons throughout the visual system. As the neural representation of sensory stimuli traverse each step of the visual hierarchy, recurrent circuits at each processing stage transform and refine it (Douglas et al., 1995; Ko et al., 2011, 2013; Cossell et al., 2015; Lee et al., 2016). Prior experimental and theoretical work in the primary visual cortex (V1) suggests that recurrent excitation amplifies responses when signals are weak in order to optimize detection (Douglas et al., 1995; Ko et al., 2011; Lien and Scanziani, 2013; Cossell et al., 2015; Lee et al., 2016), while recurrent inhibition suppresses responses when signals are strong to optimize discrimination (Anderson et al., 2000; Kapfer et al., 2007; Isaacson and Scanziani, 2011; Chettih and Harvey, 2019). Understanding what patterns of cortical activity drive either amplification or suppression is critical for a mechanistic understanding of signal transformations in the cortex. However, separating the impact of local recurrent circuits from the influence of feedforward and feedback inputs during normal physiological activity is challenging.

Past work has focused on isolating recurrent activity by removing feedforward or feedback activity. Several studies measured feedforward thalamic inputs in isolation by reversibly silencing the cortex while monitoring incoming visually evoked input using intracellular recordings (Ferster et al., 1996; Lien and Scanziani, 2013; Li et al., 2013b, 2013a; Reinhold et al., 2015). Other studies (Nassi et al., 2013; Gómez-Laberge et al., 2016) silenced higher brain areas through cooling to remove selective feedback to upstream areas and measured the changes in cortical response to a driving stimulus. In our study we take a complementary approach: we use high resolution two-photon (2P) holographic optogenetics to recreate precise experimenter-controlled patterns of neuronal activity and simultaneously measure the impact across V1 using cellular resolution 2P calcium imaging (Pégard et al., 2017; Mardinly et al., 2018). With this strategy we probe the functional logic of recurrent cortical dynamics in the absence of visual driven afferent input and unambiguously determine the causal impact of different patterns of recurrent dynamics. However, the space of 2P holographic optogenetic stimulation protocols is immense and care must be taken to parameterize a sufficiently rich, yet still feasible, probe of the recurrent circuit.

There are two main organizing principles that govern recurrent wiring in mouse primary visual cortex, particularly in Layer (L) 2/3. First, excitatory (E) and inhibitory (I) connectivity falls off with the physical distance between neurons, so that a majority of recurrent connectivity comes from neurons that are less than 200 um apart from one another (Holmgren et al., 2003; Levy and Reyes, 2012; Billeh et al., 2020; Rossi et al., 2020; Campagnola et al., 2022; Hage et al., 2022). Second, E to E connectivity is biased to occur between neurons with similar stimulus feature preferences, such as orientation tuning (Ko et al., 2011; Cossell et al., 2015; Billeh et al., 2020; Rossi et al., 2020). Many models of cortical circuits, both mathematical and conceptual, consider either spatial or feature dependent wiring, but few consider how their interaction determines overall network response. The difficulty rests in the large number circuit parameters that are needed to construct such a combined circuit model. Further, while knowledge of monosynaptic connectivity is essential for any predictive model of recurrent cortical dynamics, it is not sufficient. Both cortical nonlinearities and multi-synaptic paths complicate the relationship between physical synaptic connections and their functional influence on a specific pattern of neural activity. The governing hypothesis of our study is that by designing our 2P optogenetic stimulation protocol to probe the recurrent circuitry defined by both physical space and feature preference we will uncover the rules by which recurrence promotes either the recruitment or suppression of cortical activity.

Recent work using targeted photostimulation in the presence of a visual stimulus probed the functional ‘influence’ (Chettih and Harvey, 2019) of putative single neuron perturbation in V1. They found that co-tuned neurons tended to suppress rather than excite each other, contrary to a simple prediction from the enriched ‘like-to-like’ connectivity between excitatory neurons (Ko et al., 2011). Previous rate-base modeling work showed that to reproduce such strong like-to-like suppression, the network must have strong and specific E to I connections (Sadeh and Clopath, 2020). Meanwhile, another computational study predicted that such results would shift and yield like-to-like activation if you adjusted the contrast level of the stimulus (Cai et al., 2020). These results highlight the importance of measuring and modeling functional interactions in the intact circuit.

Moreover, the influence of a single neuron can be quite different from that of an ensemble of neurons with coordinated activity, owing to the synaptic wiring rules and integrative properties of cortical excitatory and inhibitory neurons. Indeed, multi-cell photostimulation has revealed just how diverse functional interactions are in L2/3 (Carrillo-Reid et al., 2016; Marshel et al., 2019; Russell et al., 2019; Dalgleish et al., 2020). Consequently, creating a set of generalized organizing principles for the impact and function of L2/3 recurrent circuits has remained difficult. In particular, the known converging E to I and diverging I to E connectivity imply that ensemble stimulation of E neurons will be especially potent at recruiting inhibitory circuits that can have a widespread impact on the remaining cortical network.

To define the functional logic of recurrent cortical dynamics in L2/3 of V1 we precisely photo-stimulated ensembles of excitatory neurons with 2P holographic optogenetics. We photo-activated ensembles of cells organized along two fundamental axes of the visual representation: physical space and feature space (orientation). We found that most perturbations in L2/3 of V1 generated net suppression. However, contrary to these prior findings, our data reveal two key organizing principles that eluded prior investigation. First, L2/3 recurrent circuits amplify local co-tuned activity by breaking through a recurrent blanket of suppression, but only on a very local (< 30 μm) level. Second, only spatially dense co-tuned ensembles of L2/3 neurons drive net suppression on this local scale, whereas spatially distributed co-tuned ensembles drive net excitation. A linear rate-based computational model captured these key results, but only if we incorporated a wiring rule that combines spatial- and feature-base synaptic organization. Specifically, we find that the model requires highly local like-to-like excitatory connections and the convergence of co-tuned excitatory neurons onto local co-tuned inhibitory neurons. This combination of all-optical circuit interrogation and detailed computational modeling demonstrate that neural representations in feature space and physical space intimately interact in the visual cortex. Furthermore, they outline organizing principles for the functional impact of recurrent cortical dynamics, distinguishing the conditions for when feedback amplifies input versus when it drives competitive suppression.

## Results

To determine the role of recurrent activity in L2/3, we used holographic multiphoton optogenetics to drive small ensembles of L2/3 cells in the absence of visual stimuli thereby isolating their local network impact. We employed three dimensional scanless holographic optogenetics with temporal focusing (3D-SHOT) (Pégard et al., 2017; Mardinly et al., 2018) and leveraged highly potent ultrafast opsins (Mardinly et al., 2018; Sridharan et al., 2022) which together enable the activation of dozens of cells with near single-cell resolution and millisecond precision. We simultaneously read out the activity of both stimulated and unstimulated cells using GCaMP6s. We restricted both the GCaMP6s sensor and the ChroME opsin to excitatory neurons (see Methods for details) and both imaged and photo-stimulated in 3D to obtain read/write control over a large fraction of the L2/3 V1 excitatory network. Our general methodology involved several steps: first, we tested each opsin-expressing neuron for photosensitivity and tailored the laser power for each cell to ensure reliable activation (see Methods). Next, we imaged the visual responses of this population to orientated drifting gratings, determining each neuron’s orientation tuning online. Finally, we constructed ensembles of these neurons with varied distributions of net orientation tuning and spatial locations and targeted these cells for 2P holographic photo-stimulation (for detailed cell selection criteria, see Methods).

### Heterogenous inhibition dominates recurrent network effects

We first asked how activating a small number of opsin-expressing L2/3 excitatory cells (groups of 10 targeted cells) would generally impact overall activity in L2/3 (Fig. 1A-B). Strong activation of the holographically targeted cells confirmed the efficacy of the optogenetic approach (Fig. 1C). However, there remains the possibility that stray light could accidentally activate non-targeted cells. To mitigate the possibility of including such cells in the analysis, we developed an extensive 3D calibration (see Methods, SFig. 1) resulting in a high-quality optical point-spread function (PSF) and a physiological PSF, which defined our off target exclusion zone (see Methods, Fig. 1D). Henceforth, all neurons lying within this exclusion zone are excluded from analysis. Additionally, by using SepW1-Cre x Camk2a-tTA x tetO-GCaMP6s mice, where Cre-dependent opsin expression is intentionally sparse, we found only a minimal difference in nearby activation of opsin-negative cells outside of this exclusion zone (SFig. 2), further backing our threshold for cell exclusion.

**Figure 1.**
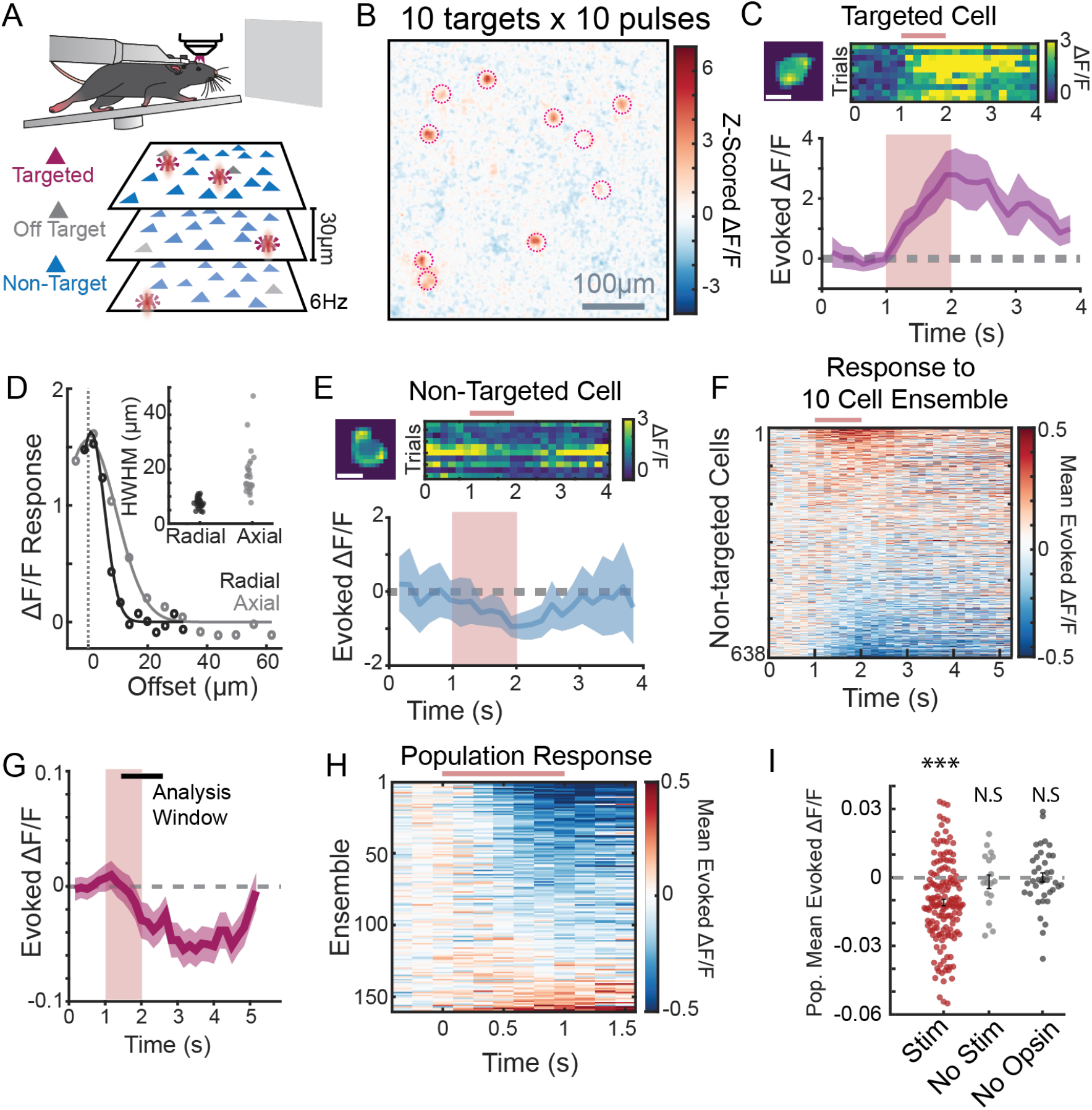
Stimulation of 10-cell ensembles recruits net inhibition: **A:** Schematic of the experimental setup. Head-fixed mice are allowed to run on a treadmill while watching a neutral grey screen. Through a cranial window, cells from three planes spaced 30 μm apart are imaged at 6 Hz. Cells from any plane can be targeted for photostimulation (magenta), cells adjacent to photostimulated cells (including offset axially) are categorized as ‘off target’ and excluded (grey), while remaining detected cells are ‘non-target’ cells and used for analysis (Blue). **B:** A representative image superimposing all three planes of imaging during stimulation of a representative 10 cell ensemble. Image pixels are z-scored over the entire recording and averaged by trial type. **C**: Response of a Targeted Cell to photostimulation. Top Left: Image of the cell used in analyses (aka cell mask), scale bar 10 μm. Top Right: heatmap of 10 photostimulation trials, stimulation time noted by maroon bar. Bottom: Mean ± 95% confidence interval (c.i.) Evoked ΔF/F from that targeted cell in response to stimulation. Maroon box denotes stimulation time (10 Pulses and 10 Hz). **D:** Representative physiological point spread function (PPSF) radially (black) and axially (grey), aligned to peak (dotted line). Inset PPSFs from 25 cells throughout the field of view. **E:** Response of a Non-Target cell to ensemble stimulation, as in C. **F:** Mean response of 638 non-target cells from a single FOV in response to a representative ensemble stimulation. Cells are sorted based on their response magnitude. Stimulation time noted by maroon bar. **G:** Mean ± 95% c.i. of all 638 non-target Cells from F. Maroon box indicates stimulation time, black bar indicates analysis window. **H:** Population response, i.e., mean response of all non-target cells, in a field of view to 160 unique 10 cell ensembles. N=160 Ensembles, 18 FOVs, 13 Mice. **I:** Mean population response for each ensemble stimulation (maroon, n=160 Ensembles, 18 FOVs, 13 Mice), no stimulation controls (grey, n=18 FOVs, 13 Mice), or no opsin controls (black, n=38 ensemble, 1 FOV, 1 Mouse). Mean ± SEM of condition in black. *** significantly different from 0, N.S. not significant (Stim p<1.7e-8, No Stim p=0.79, No Opsin p=0.96; signed rank test)

Non-targeted neurons displayed a variety of effects, with individual cells responding distinctly to different ensembles (SFig. 3). However, the majority of cells were suppressed in response to ensemble stimulation (Fig. 1E-G, SFig. 3B). Across a large set of such experiments (160 unique 10-cell ensembles in 18 fields of views (FOVs) in 13 different mice) we found that photo-stimulation suppressed mean population activity (mean effect: −0.011±0.0014 ΔF/F, p<1e-10, Wilcoxon rank sum test, Fig. 1H-I). These results indicate that photo-stimulation of a small number of L2/3 excitatory neurons recruits net inhibition across the entire population.

However, ample evidence suggests that different patterns of neural activity can recruit distinct recurrent circuits that might preferentially drive either activation or suppression of activity (Chettih and Harvey, 2019; Marshel et al., 2019). Thus, we next asked whether the overall sign and magnitude of recurrent activity during holographic photo-stimulation depended on how the pulses were added to the system. First, we varied the total number of pulses delivered to a group of 10 targeted neurons between 1 and 50 pulses per cell (~10 to ~500 total pulses added), while holding the pulse frequency and ensemble size constant (10 Hz and 10 cells, respectively). We observed net suppression across all these conditions, with a monotonic increase in suppression with increased number of spikes (p<1.7e-8, ANOVA, N=76 Ensembles, 5 FOVs, 2 Mice, SFig. 4A). Next, we varied the stimulation frequency, while holding ensemble size and total added pulse number constant. In contrast to the previous result, varying the rate of stimulation did not change the magnitude of the net mean suppression (p>0.74 Anova N=46 Ensembles, 2 FOVs, 2 Mice, SFig. 4B). Likewise, varying ensemble size (3-33 cells) while adding a fixed total number of spikes also drove net suppression that did not vary (mean ΔF/F: −0.0045 ± 0.0027 33 pulses in 3 cells, −0.0081 ± 0.0023 10 pulses in 10 cells, −0.0094 ± 0.0031 3 pulses in 33 cells, p >0.78 Anova N=99 Ensembles, 11 FOVs, 5 Mice, SFig. 4C). These results demonstrate that the primary driver of network suppression is the total number of added spikes, not the frequency of stimulation nor the size of the ensemble.

Based on these results, we focused on the effects of adding 100 total pulses to 10 targeted neurons, which represents a modest perturbation to the system that still drove reliable and readily quantifiable effects. Importantly, such modest perturbations are well captured through simulations of associated network models (see below) since they can be modeled as a linear perturbation around the network’s steady state.

While it is true that ensemble stimulation leads to suppression *on average* across the population, there remains significant heterogeneity of response, with a significant number of neurons showing activation rather than suppression. Specifically, we found 2.34 ± 0.09% of non-targeted cells were significantly activated (i.e., 99% CI excludes 0, false discovery rate 1%) and 5.58 ± 0.19% of non-targeted cells were significantly suppressed (SFig. 3B-C). The central goal of this study is to explain this heterogeneity based on the joint physical and feature space properties of the neurons both in the stimulated ensemble and recorded populations.

### Cortical space organizes the impact of recurrent dynamics

Physical space in the sensory neocortex represents a fundamental axis of circuit organization owing to both the topographic mapping of sensory inputs onto cortical tissue and the anatomy of cortical neurons (Dräger, 1975; Wagor et al., 1980; Garrett et al., 2014; Zhuang et al., 2017). Thus, we hypothesized that the sign, scale, and magnitude of recurrent circuit influence might vary substantially with distance from the targeted ensemble. To test this, we quantified the impact of ensemble photo-stimulation as a function of distance from each targeted location and ensemble (see Methods). Indeed, we found that despite the overall mean suppression described above, cells proximal to the stimulated ensemble but outside of the off target exclusion zone were reliably activated, while cells further away from a target were suppressed (< 30 μm from a target mean ΔF/F: 0.044 ± 0.005, p<1.1e-10; 50-150 μm from a target mean ΔF/F: −0.013 ± 0.001, p<4.0e-17, Signed Rank Test; Fig. 2A). Beyond that distance, the sign of the modulation stayed negative and slowly returned to zero as the distance increased.

**Figure 2:**
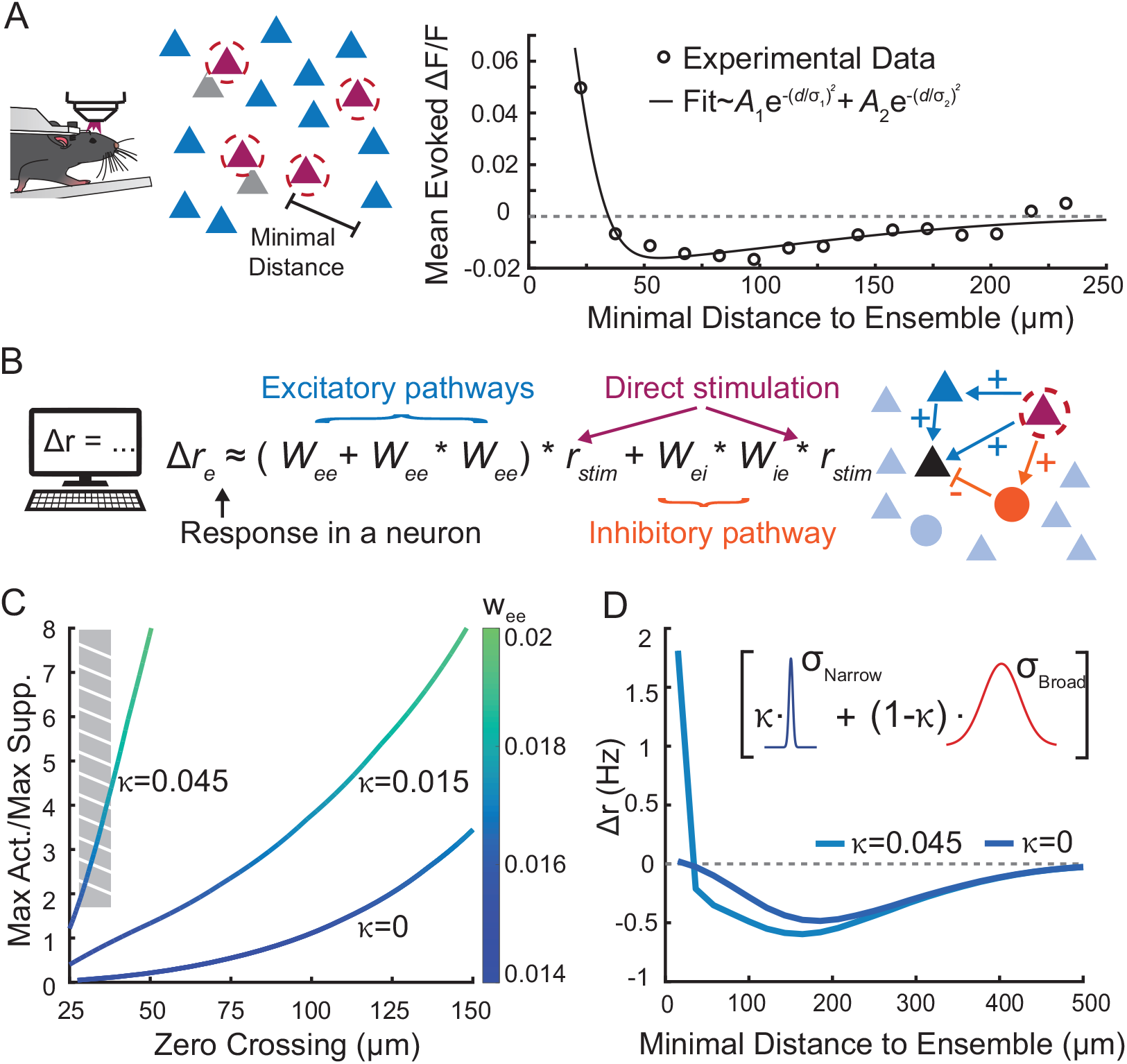
Decomposing population responses reveals nearby activation and surround suppression. **A:** Left: Schematic of the minimal distance metric for non-target cells (magenta: targeted cells, gray: off-target cells, blue: non-targeted cells). Right: Non-targeted cell responses to optogenetic stimulation as a function of minimal distance to ensemble (open circles) fitted to a sum of Gaussian spatial functions (A_1_ = 0.1956, σ_1_ = 22.07 μm, A_2_ = −0.021, σ_2_ = 147.31 μm). Cells near to the stimulated ensemble were reliably activated, while cells further away from a target were suppressed (<30μm from a target mean ΔF/F: 0.044 ± 0.005, p<1.1e-10; 50-150 μm from a target mean ΔF/F: −0.013 ± 0.001, p<4.0e-17, Signed Rank Test). Bin sizes are 15 μm. **B:** Schematic of all monosynaptic and disynaptic pathways resulting in a change in the baseline firing rate (blue: excitatory pathways, orange: inhibitory pathways). * denotes convolution in space. **C:** The zero crossing and relative strength of nearby excitation (max activation / max suppression) as a function of recurrent excitatory strength (*w_ee_*) and biased connections on the narrow spatial scale (*κ*). The gray stripe box indicates the experimentally observed data regime and illustrates the need of the additional spatial to capture the nearby excitation observed in the data. **D:** Non-targeted cell responses in the model as a function of minimal distance to ensemble for different values of *κ*. Inset: schematic show the narrow vs. broad spatial scales in the model (see Methods for more details).

This spatial pattern of nearby excitation and surround inhibition characterizes the spatial response function of a minimal recurrent circuit and has often been considered as a basis for lateral competition in the cortex (Cavanaugh et al., 2002; Derdikman et al., 2003; Kang et al., 2003; Isaacson and Scanziani, 2011; Adesnik et al., 2012; Shushruth et al., 2012). Phenomenologically, it can be captured as the difference of a narrow excitatory and a broader inhibitory gaussian spatial function (Fig. 2A, solid line; excitatory spread: 22 μm, inhibitory spread: 147μm). Moreover, these experiments demonstrate that activation of even a small number of L2/3 excitatory neurons is sufficient to generate this ‘check-mark’ shape. However, exact mechanistic circuit basis giving raise to this shape remains unclear.

To address this question, we developed a computational model of these targeted optogenetic perturbations (see Methods). Our goal was to understand how connectivity principles in L2/3 recurrent circuits could explain why recurrent activation is extremely local but why suppression dominates at larger distances from the stimulated ensemble. We wired the simulated circuit based on previously acquired connectivity data (Rossi et al., 2020; SFig. 5A-D), and modeled the dynamics of the population with a two-dimensional neural field model (Huang et al., 2019, see Methods). Due to the modest size and strength of the optogenetic perturbations we considered the network response as linear perturbations around a steady state firing rate solution. Further, because the experiments were performed in absence of a visual input, we could assume that the neurons in the network have a low gain response, implying that the effective connectivity strength is relatively weak. This allowed us to investigate the network perturbation via a synaptic pathway expansion with just a few terms (Ocker et al., 2017; Sadeh and Clopath, 2020). Specifically, we considered monosynaptic and disynaptic excitatory connections (E→E and E→E→E) and disynaptic inhibitory connections (E→I→E) (Fig. 2B).

After fitting the spatial components of the model (see Methods), we are left with two free parameters that correspond to the strengths of these pathways: *w_ee_* (the effective strength of E→E connections) and *w_eie_* (the effective strength of the inhibitory pathway). We find that nearby excitation and surround inhibition arises for a variety of parameter values, with near identical shapes arising for fixed values of *w_ee_/w_eie_*. After fixing *w_eie_* and varying *w_ee_* for simplicity, we find that we can adjust both the zero crossing of this curve and the strength of the nearby excitation (Fig. 2C). However, we see that this model is unable to pass through the experimentally observed data regime (Fig. 2C, gray box, SFig. 5E, see Methods). Specifically, when these parameters are adjusted to match the experimental observed cross at ~35 μm, the model fails to capture the relative strength of nearby excitation to more distant suppression. To capture this key detail, we reasoned that we needed to add an additional spatial scale to the model (Fig. 2D). Based on recent work (Kwan and Dan, 2012; Kondo et al., 2016; Ringach et al., 2016; Yu et al., 2020), L2/3 neurons in mouse V1 have unique connectivity rules on a narrow spatial scale of (< 50 μm) leading to small columnar structures that is not strictly a salt-and-pepper organization. Adding in such a tight spatial component (i.e., *κ* >0, see Methods) allowed the model to simultaneously capture both the nearby excitation and the appropriate zero crossing between excitation and suppression.

Similar to how different sensory stimuli will drive different spatial distributions of activity, we next asked how the spatial distribution of targeted cells would impact either the suppression or activation of recurrent activity. To investigate this question experimentally, we activated ensembles of 10 neurons that were either distributed or clustered in space (see Fig. 3A). Indeed, we found that activating a spatially compact ensemble drives much more surround inhibition than stimulating a spatially diffuse ensemble (linear regression of mean ΔF/F vs spread slope: 1.3e-4 ΔF/F per μm spread, p<1.4e-5; Fig. 3A). In contrast, the spread of the ensemble did not alter nearby excitation (linear regression of mean ΔF/F vs spread slope: 3.5e- 5, p=0.57; SFig. 6A-C). More precisely, and in line with our computational model, the level of surround inhibition (at 50-150 μm) increased as the spatial distribution of the ensemble’s component neurons decreased (Fig. 3B). To understand what accounts for this, we examined in the model the relative strength of the E→E and E→I→E pathways as a function of the spatial spread of the ensemble. We found that as the ensemble spread decreases, both synaptic pathways increase in magnitude, but the strength of inhibitory pathway increases faster, leading to the observed effect (Fig. 3C; slope is ~9% greater for the inhibitory pathway).

**Figure 3:**
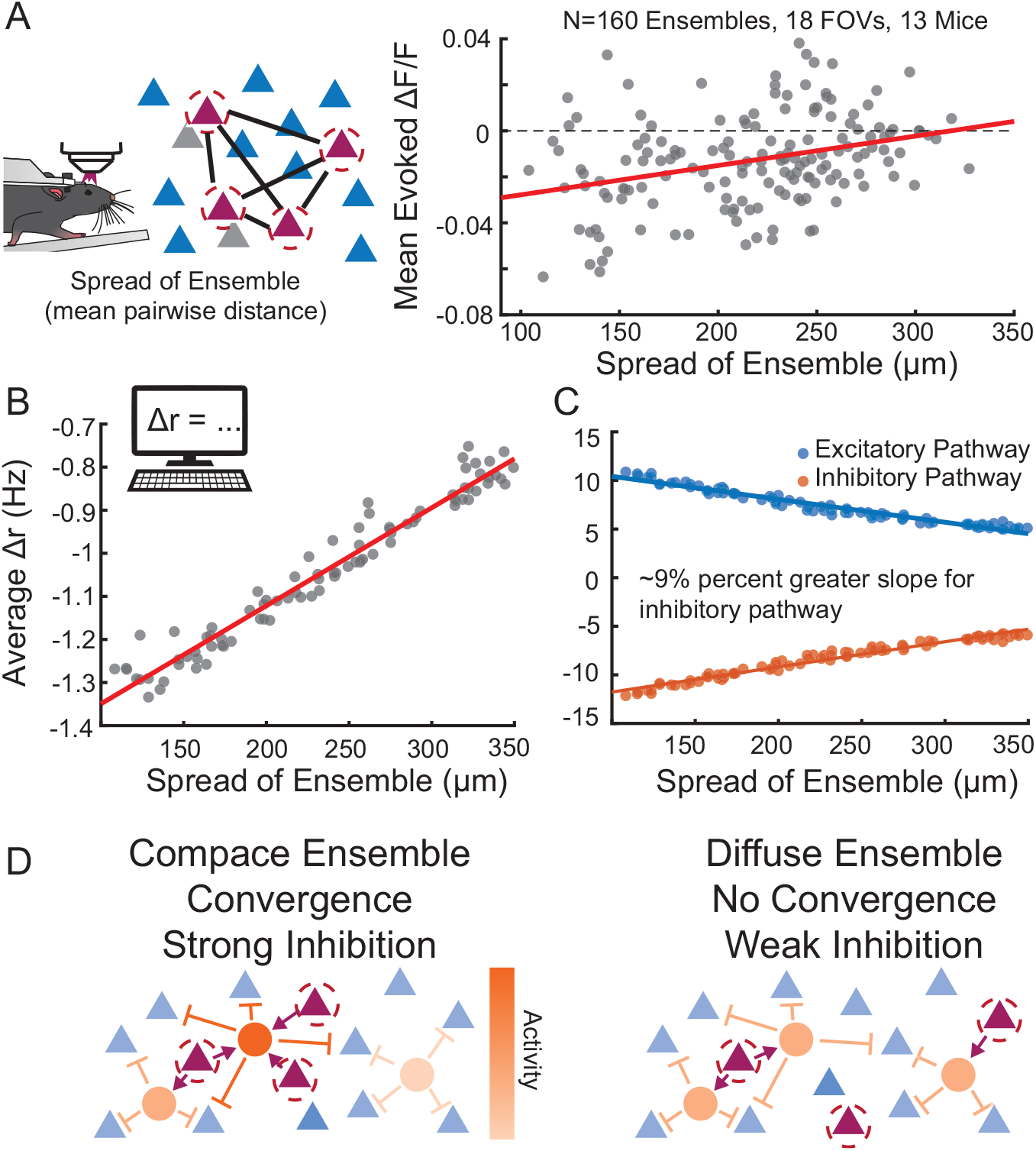
Tight ensembles recruit stronger network suppression because of converging inhibitory pathways. **A:** Left: Schematic of ensemble spread metric (mean pairwise distance). Right: Non-targeted cell responses averaged across the population (N=160) for ensembles with different spreads. Activating a spatially compact ensemble drives more surround inhibition than stimulating a spatially diffuse ensemble (linear regression of mean ΔF/F vs spread slope: 1.3e-4 ΔF/F per μm spread, p<1.4e-5). **B:** Same as A except for the network model. **C:** Strength of the model excitatory (blue) and inhibitory (red) pathways as a function of ensemble spread, showing that as the ensemble spread decreases, the inhibitory pathway shows a greater level of recruitment. **D:** Schematics illustrating the ability of the targeted cells to activate inhibitory pathways via convergences for compact (left) and diffuse (right) ensembles.

These findings are consistent with the idea that strong inhibition derives from the convergence of excitatory activity onto individual inhibitory neurons. In the case of a spatially compact ensemble individual inhibitory neurons receive input from multiple directly stimulated cells. These super activated inhibitory cells then feedback divergently inhibiting the entire network (Fig 3D).

### Feature space organization of the impact of recurrent dynamics

In addition to physical space, feature space represents a second axis of the functional organization of cortical circuits. In mouse V1, orientation-tuning is a crucial feature that is not structured in physical space, unlike the orientation columns and pinwheels of monkeys, cats, and other species (Ohki et al., 2005; Ohki and Reid, 2007; Bonin et al., 2011). Despite the lack of local organization, feature space is known to influence both synaptic connectivity and the functional influence of individual neurons in mouse V1 (Ko et al., 2011; Cossell et al., 2015; Chettih and Harvey, 2019). However, as most reports focus on the impact of individual neurons, it is unknown if multiple neurons defined in feature space synergize to drive recurrent activity. One hypothesis is that a co-tuned (i.e., iso-oriented) group of excitatory neurons, analogous to spatial clustering in orientation space, should drive strong network activity due to convergent excitation onto the same postsynaptic excitatory cells (Marshel et al., 2019).

To test this hypothesis, we presented mice with a randomized series of full screen drifting gratings each trial presenting one of 8 cardinal directions of motion and calculated tuning curves for each neuron online, generating ensembles of cells varying in preferred-orientation and orientation selectivity. We summarized the selectivity of an ensemble using an ‘ensemble OSI’, i.e., the OSI of the average of the tuning curves (see Methods, Fig. 4A-B). To optimally select exemplar ensembles (among the ~10^23^ possible ensembles – 1,000 choose 10) we created a discrete optimizer (see Methods) that designs distinct ensembles by automatically choosing eligible cells that fall within a targeted spatial and OSI range. We used this optimizer to ensure that the chosen ensembles were evenly distributed across feature (orientation) space. Surprisingly, we found that ensembles with higher ensemble OSI did not generate greater network effects when averaged across all non-targeted neurons (SFig. 6D-F, no significant correlation between ensemble OSI vs the total population response, p = 0.26 linear regression; or between ensemble OSI and nearby, mid distance, or far cells’ responses; p=0.084, p=0.27, and p=0.89 respectively by linear regression).

**Figure 4:**
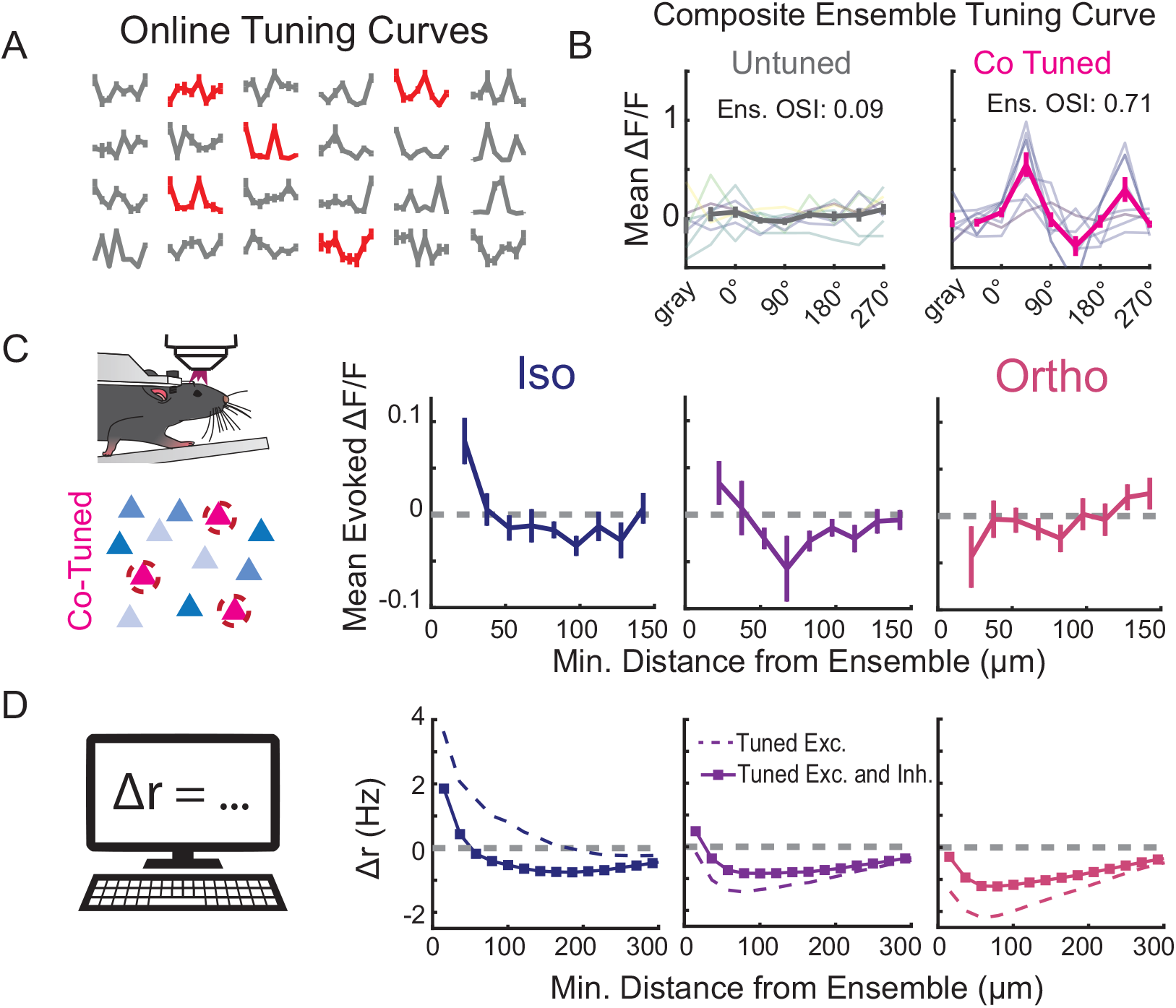
Relative tuning of non-targeted cells determines response to stimmed ensembles. **A:** Representative online tuning curves from 24 non-targeted cells. Each point of each curve is the mean ± SEM normalized Ca response to a drifting visual grating. Red cells indicate cells that were selected for an individual ensemble. **B:** Two representative composite ensemble tuning curves, i.e., the mean of the tuning curves of the 10 cells that make up an ensemble, are shown. Left a representative untuned ensemble (low ens. OSI), right a representative co-tuned ensemble (high ens. OSI). Individual tuning curves lighter colors, mean ± SEM dark bold colors. **C:** Left: Schematic showing a co-tuned ensemble. Right: Nontargeted cell responses ± SEM as a function of their minimal distance to the ensemble according to their relative tuning with the stimmed ensemble (Left: Δθ = 0° (Iso), Middle: Δθ = ±45°, Right: Δθ = 90° (Ortho)). The included cells were responsive to visual stimuli (p-value < 0.05) and tuned (OSI > 0.25). Nearby iso-oriented cells (< 30 μm) are activated dramatically more than those cells that prefer orthogonal stimuli (p<0.0035, Wilcoxon one-sided ranked sum test, N=17 Ensembles, 8 FOVs, 3 Mice). Bin sizes are 15 μm. **D:** Same as C, except for the network model with (solid) and without (dashed) tuned E→I connections.

Based on the principle of selective like-to-like connectivity, we next asked whether cotuned ensembles might preferentially impact non-targeted neurons that share feature preference. To test this hypothesis, we restricted our analysis to the ‘co-tuned ensembles’ and divided the non-stimulated cells based on their relative preferred orientation. Such a ‘co-tuned ensemble’ consisted of individually tuned members (mean OSIs of ensemble members>0.5) with similar tuning preferences (ensemble OSI >0.7). We find that nearby cells (<30μm) that also prefer the same orientation as the stimulated ensemble (i.e., iso-oriented cells), are activated dramatically more than those cells that prefer orthogonal stimuli (p<0.0035, Wilcoxon one-sided ranked sum test, N=17 Ensembles, 8 FOVs, 3 Mice, Fig. 4C). Strikingly, these nearby orthogonally oriented cells were instead highly suppressed. These results demonstrate that the organization of an ensemble in feature space – in this case orientation preference – profoundly influences its recurrent impact on specific cells in the cortical network.

To explain this switch from like-to-like excitation to like-to-unlike suppression, we used our computational model to determine which features of circuit connectivity are required to generate it. We considered three hypotheses: (1) the orientation dependence of recurrent inputs could emerge on their own based purely spatial connectivity rules and salt-and-pepper orientation tuning, (2) like-to-like E→E connectivity but random E→I and I→E connectivity could explain it, or (3) orientation specificity would be required in all of these pathways. In line with prior work, we assumed that the orientation preferences of individual neurons are inherited from feedforward projections and not spatially organized.

When synaptic connectivity only followed a spatial wiring rule with no specificity in orientation space, we found no difference in the recruited recurrent activity of iso-oriented vs. orthogonally oriented neurons (SFig. 7), demonstrating that pure spatial rules are not sufficient on their own to explain the experimentally observed recurrent dynamics. Adding in like-to-like connectivity between excitatory neurons (Ko et al., 2013; Rossi et al., 2020) reproduced orientation-preference dependent effects, qualitatively similar to the experiment results (Fig. 4D; dashed). Specifically, cells that were iso-oriented to the photo-stimulated tuned ensemble showed excitation, while those that are orthogonally orientated showed suppression. However, the model with this wiring scheme substantially overestimated iso-oriented excitation and orthogonally oriented suppression at all distances, and completely failed to capture the iso-oriented surround inhibition beyond 50 μm. Finally, when the model incorporated like-to-like excitatory-to-inhibitory and inhibitory-to-excitatory connections, as recently suggested by (Znamenskiy et al., 2018), it accurately reproduced the experimental data both qualitatively and quantitatively (Fig. 4D; solid). These results imply that feature-specific synaptic connectivity across all three synaptic pathways is essential to explain the space and feature-dependence of recurrent cortical dynamics.

### Interactions between physical space and feature space

Thus far, we have only considered how the geometric distribution (Fig. 3) and feature preferences of an ensemble (Fig. 4) govern cortical recurrent dynamics independently. However, since sensory stimuli will necessarily recruit recurrent dynamics that vary jointly across these two dimensions, we hypothesized that the spatial distribution and feature preference of an ensemble should interact to determine the resultant impact of ensemble photo-stimulation on the cortical network. To investigate this hypothesis, we used the discrete optimizer to identify co-tuned or untuned ensembles that were either spatially compact or spatially distributed and photo-stimulated them while observing the activity of the non-target cells. First, we found that the spatial spread of an un-tuned ensemble (ensemble OSI <0.3 and mean OSIs of ensemble members<0.5, see Methods) did not affect its net recurrent impact, such that for both spatially compact and diffuse ensembles we observed the characteristic nearby excitation and surround inhibition when computed across all non-targeted neurons (Fig. 5A, grey traces). However, the spatial spread of a co-tuned ensemble profoundly influenced its recurrent effects: compact, co-tuned ensembles generated no nearby excitation and instead showed nearby suppression, whereas a spatially diffuse co-tuned ensemble generated the more typical center/surround effects (Fig. 5A, pink and purple traces; nearby activity co-tuned, close ensemble (N = 8) vs. co-tuned far ensemble (N=17) p <0.008, Wilcoxon one-sided ranked sum test).

**Figure 5:**
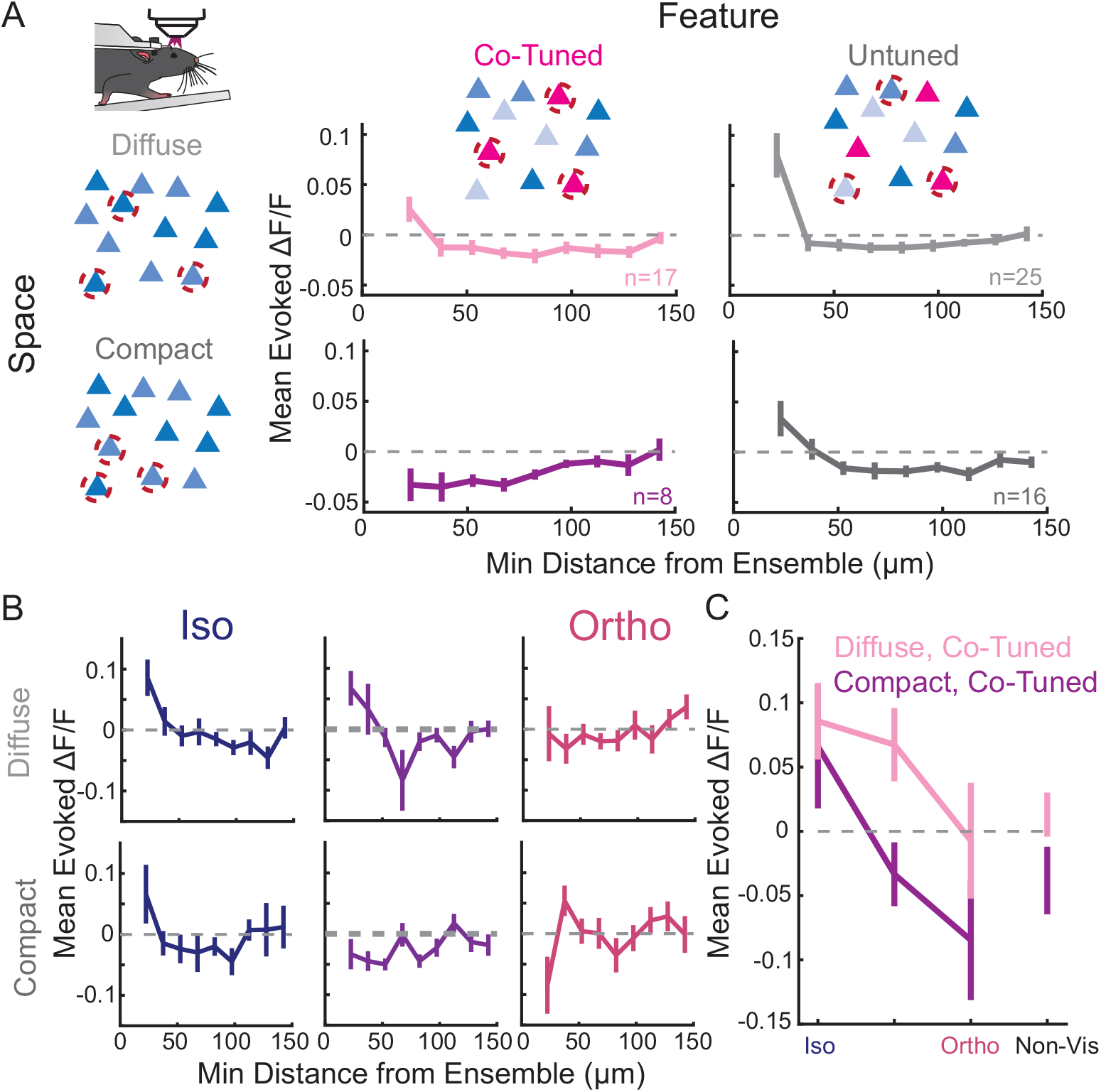
Space and feature properties of the stimmed ensemble shape the population responses. **A:** Ensembles are divided by two categories, whether they are diffuse (mean distance > 200 μm, top) or compact (< 200 μm, bottom) vs cotuned (Ens. OSI > 0.7, Left) or Untuned (< 0.3, Right). Non-targeted cell responses ± SEM as a function of their minimal distance to the ensemble. **B:** Co-tuned ensembles divided by mean separation (as in A), but non-targeted cells are separated by relative tuning with the stimmed ensemble (Left: Δθ = 0° (Iso), Middle: Δθ = ±45°, Right: Δθ = 90° (Ortho)). **C:** Non-targeted cell responses from the first bin of panel B as a function of their relative tuning with the stimmed cotuned ensembles.

This result appears at odds with previous optogenetic results, which have suggested that a co-tuned ensemble should evoke a population response reminiscent of a visual image containing a drifting grating (Marshel et al., 2019). To reconcile these results, we asked how the activity of visually responsive cells varied across spatially diffuse and compact co-tuned ensembles, as a function of their orientation preference (Fig. 5B-C). We observed that diffuse co-tuned ensembles recruited very little nearby suppression in neurons at any orientation. In contrast, compact co-tuned ensembles, activated nearby iso-oriented cells, while nearby cells at other orientations were reliably suppressed. Further, non-visually responsive cells were also suppressed during the stimulation of such ensembles. In total, these data suggest that compact, co-tuned ensembles were able to sharpen the input signal locally, activating iso-oriented cells while suppressing all other cells. Diffuse co-tuned ensembles, however, lacked this ability to significantly suppress nearby ‘unlike’ cells.

To unpack these results further, we turn again to our computational model, which now consists of both the spatial- and feature-based wiring rules necessary to reproduce our core experimental findings. Indeed, simulations showed that a diffuse, co-tuned ensemble generated effects that are similar to previous results: nearby excitation and suppression inhibition (Fig. 6A; pink curve), while compact, co-tuned ensembles drove suppression across all distances (Fig. 6A; magenta curve). Decomposing this result into the direct excitatory and disynaptic inhibitory pathways, we found that while the compact ensemble recruits both more inhibition and more excitation than a diffuse ensemble (Fig. 6B), however, inhibition tends to dominate the net effects on firing rates. This observation illustrates an interesting tradeoff between the E→I→E inhibitory pathway and the excitatory pathways as one compresses the cotuned ensemble. Namely, nearby suppression replaces nearby excitation as the ensemble shrinks in space.

**Figure 6:**
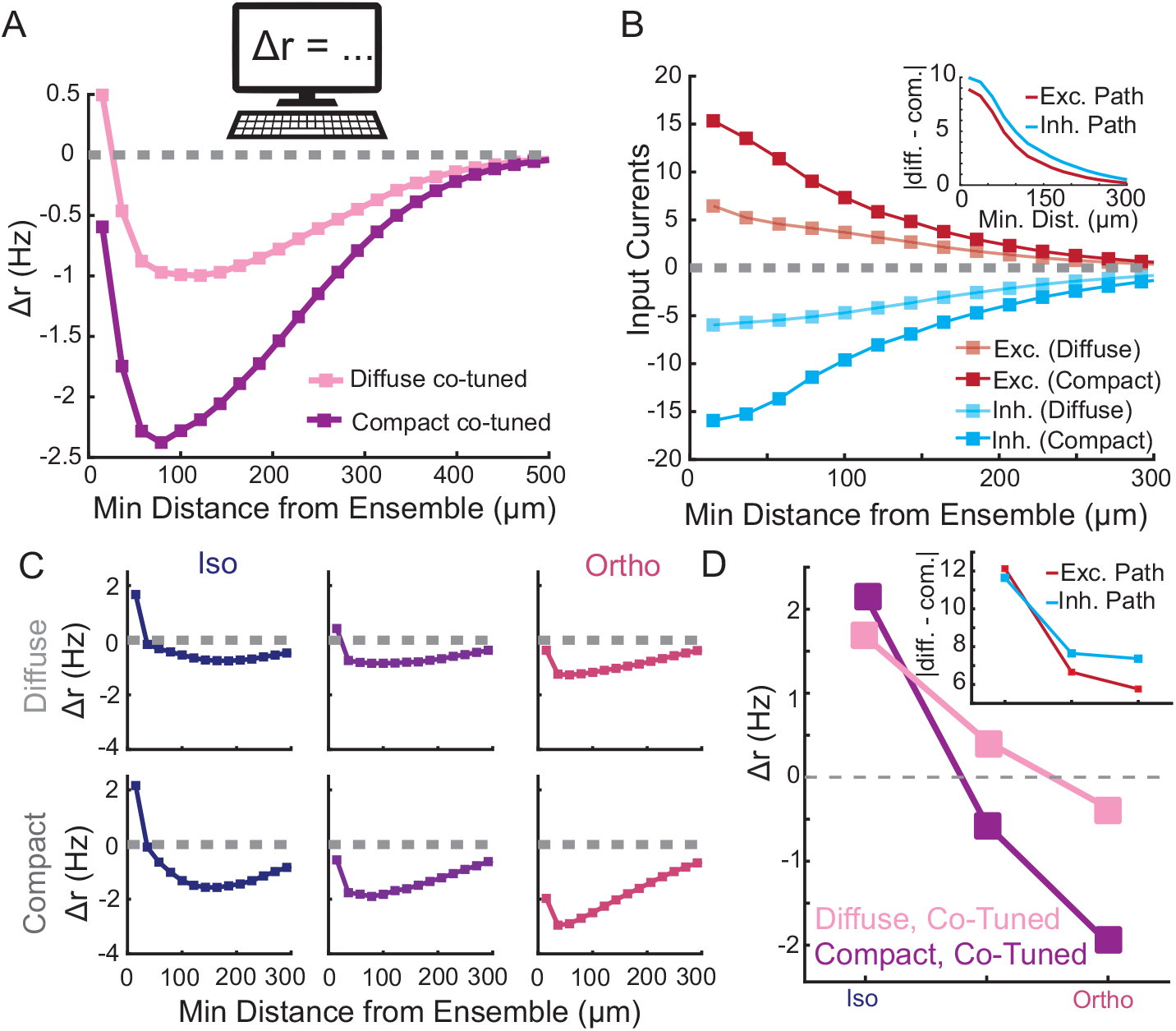
Network model with both spatial- and feature-based tuned connections wiring rules can recapitulate core experimental results. **A:** Non-targeted cell responses in the network model as a function of their minimal distance to the ensemble according to the ensemble spread and tuning. **B:** The excitatory (red) and inhibitory (blue) input pathways for the spatially diffuse and compact co-tuned ensembles. Inset: Absolute difference of excitatory and inhibitory paths showing that while both pathways increase in magnitude for compact ensembles, the inhibitory pathway shows a larger increase. **C:** Non-targeted cell responses in the network model as a function of their minimal distance to the ensemble according to their relative tuning with the stimmed co-tuned ensembles. **D:** Non-targeted cell responses from the first bin of panel C as a function of their relative tuning with the stimmed co-tuned ensembles. Inset: Absolute difference of excitatory and inhibitory pathways.

After decomposing these effects based on the orientation preference of the non-targeted neurons, we find that the model continues to agree with our experimental results: compact, co-tuned ensembles sharpen the input signal locally more so than diffuse, co-tuned ensembles (Fig. 6C and 6D). Again, we can decompose this result into the two pathways, and again we that it is the stronger recruitment of the inhibitory pathway, in this case via cells at non-iso orientations, that lead to this result (Fig. 6D, inset).

## Discussion

Recurrent activity in V1 could serve to amplify sensory input when signals are weak (Douglas et al., 1995; Ferster et al., 1996; Ichida et al., 2007; Ko et al., 2011; Lien and Scanziani, 2013; Li et al., 2013b; Cossell et al., 2015; Lee et al., 2016), but drive competition among stimuli when signals are strong (Anderson et al., 2000; Kapfer et al., 2007; Isaacson and Scanziani, 2011; Chettih and Harvey, 2019; Mossing et al., 2021). However, as definitively isolating the role of recurrent circuits from the impact of feedforward or feedback input has not been feasible, fundamental concepts on the role of recurrent activity in cortical computation remain untested. By selectively activating small ensembles of L2/3 cells in the absence of visual input we could isolate the role of local recurrent activity using multiphoton holographic optogenetics and calcium imaging. By leveraging our ability to design unique ensembles of L2/3 neurons whose properties varied across both physical space and feature space, we systemically tested the role these two fundamental axes have in driving recurrent activity. Moreover, by combining this experimental approach with detailed computational modeling, we proposed a wiring rule that depends on both spatial and feature properties of cells and investigated the implications of such a rule.

We found that recurrent circuits could either amplify or suppress cortical activity depending on the spatial distribution and tuning of the presynaptic ensemble and the location and tuning of the postsynaptic cells. The vast majority of photo-stimulation patterns led to net suppression of cortical activity, but the sign, scale and magnitude of recurrent network modulation followed a specific logic. For example, we found that compact, co-tuned ensembles largely drive suppression, while spatially distributed co-tuned neurons drive amplification, but only of nearby neurons, otherwise they likewise drive suppression. Taken together, our results demonstrate that the recurrent circuitry defined by feature space and physical space jointly determine the impact of recurrent circuits.

Importantly, we used relatively modest perturbations (~10 action potentials added to ~10 neurons) to avoid pushing the system out of its physiological operating range. Further, from a computational perspective, modest perturbations are easier to interpret since they evoke small deviations from the steady-state firing rate solution. Indeed, we were able to construct network simulations with few parameters that could accurately capture our initial experimental results and qualitatively predict further experimental outcomes. Investigation of the model suggested that like-to-like connectivity not only between excitatory neurons, but also between excitatory and inhibitory neurons, was important for explaining the results. A narrow spatial scale of like-to-like E→E connectivity, as suggested previously (Kwan and Dan, 2012; Ringach et al., 2016; Yu et al., 2020), was likewise essential for accurately predicting the experimental data.

Several recent studies have also use 2P optogenetics to probe functional connectivity in mouse visual cortex. One recent study (Marshel et al., 2019) found that optogenetic stimulation of tuned ensembles in mouse V1 primarily generated recurrent amplification of co-tuned activity. Under most conditions in our data, suppression dominates, with recurrent excitation confined to a close distance to targeted cells. One possible explanation for this apparent discrepancy is that we deliberately made smaller perturbations of co-tuned ensembles limited to about ten neurons, while this other study aimed to recruit larger numbers of neurons and possibly at higher firing rates. It is possible that increasing the number of photo-stimulated neurons in the co-tuned ensemble in our experiments would increase excitation relative to inhibition which could lead to net amplification. However, more generally, since we found that net impact of recurrent circuits depends on several key properties of the neural activity pattern, it is also possible that if this other study took these factors into account our results could be in more agreement.

Two recent studies in mouse V1 either targeted single (Chettih and Harvey, 2019) or multiple (Russell et al., 2019) neurons in L2/3 with 2P photostimulation largely generated suppression across the network, but also some nearby excitation. While this appears to better match our results, these studies also observed much more substantial like-to-like suppression, leading one of the studies to hypothesize that recurrent networks are primed for competition rather than amplification (Chettih and Harvey, 2019). However, there are two key differences between this study’s approach our the one we used here. First, they targeted single neurons rather than ensembles (Chettih and Harvey, 2019). Since most synapses in cortex are weak, the impact of adding spikes to one neuron could be substantially different than when activating ten. For example, cortical somatostatin cells, which are a major source of recurrent inhibition in L2/3, are only recruited effectively by the activation of two or more L2/3 pyramidal cells (Kapfer et al., 2007). Additionally, the notion of compact versus diffuse nature of an ensemble, which we found to be a critical determinant of net impact, has no meaning for single-neuron perturbations and was not explored in Russell et al. (2019).

Furthermore, both aforementioned studies conducted their experiments while presenting visual stimuli to the animal. This may have been necessary to make it possible to measure the very small effects of single-neuron photo-stimulation (Chettih and Harvey, 2019) or influence behavioral performance (Russell et al., 2019), yet it also means the network was in a state that was potentially dominated by non-recurrent sources of input. This could shift the dynamical state of network, and potentially lead to a stronger recruitment of suppression. Finally, the magnitude and sign of the like-to-like modulation was not computed across different distances from the optogenetically targeted neuron (Chettih and Harvey, 2019). Since we also found that this distance was a key modulator of like-to-like effects, it is possible that such analysis might reveal better agreement with our data.

Perhaps one key issue with comparing results of 2P optogenetic targeting across studies is the enormous potential parameter space of the perturbations. These include the number, spacing and featuring tuning of the targeted neurons, but also the precise temporal pattern of optically evoked activity and total number of spikes. Some of these are readily under user-control (such as the spacing of the targeted neurons, the number of pulses delivered to each neuron) and some are often not, such as the exact number of neurons that are actually photo-stimulated and the temporal pattern of their activity. Additionally, the optical approach and the microbial opsin used can profoundly influence the pattern and magnitude of activity. Standardizing these parameters should aid in better comparison. More generally, using an approach that ensures specific numbers and temporal patterns of the evoked spikes, as recently described, should obviate the need for matching these parameters.

With respect to our computational modeling, a similar study (Sadeh and Clopath, 2020) also made use of a linear rate-based model to explore the pathways driving recurrent circuit impacts during photo-stimulation of a small number of neurons. Like this study, we also found that the E→I pathway must be sufficiently strong and feature-specific to explain the large amount of suppression observed. However, this previous study largely focused on explaining how the optogenetic perturbation of a single cell influences recurrent activity. Here, we were able to further develop a model that simultaneously incorporates space- and feature-based wiring rules due to the larger number of neurons in the stimulated ensemble. Specifically, by exploring different spatial distributions of the activated ensemble, we observed interesting trade-offs between like-to-like amplification and suppression on different spatial scales. Such ensemble geometries are simply not possible in single-cell perturbation experiments. While another computational study (Cai et al., 2020) also examined the effects of stimulating a larger number of neurons, they only considered co-tuned ensembles and did not vary their spatial distribution.

Our combined *in vivo* and *in silico* interrogation of recurrent dynamics helps define an elementary logic for the impact of recurrent circuits on cortical activity. Our data and analysis demonstrate that recurrent activity does not simply amplify or suppress activity. Rather, the impact of recurrent circuits in L2/3 of mouse V1 depends jointly on physical space and feature space, both from the perspective of the activated ensemble and the other neurons in the network. Our computational modeling points to clear and testable predictions for the underlying circuitry that would support these local computations, which may be borne out in future experiments. The richness in recurrent modulation we discovered here might conceivably be suited to matching the sophisticated demands on the processing of complex images, such as occur naturally in the world. More generally, these principles our work reveals may constitute an elemental neural syntax of cortical transformations by recurrent circuits.

## Methods

All experiments were performed in accordance with the guidelines and regulations of the ACUC of the University of California, Berkeley. Protocol #AUP-2014-10-6832-2.

### Mice

All calcium imaging experiments were performed in mice of both sexes expressing GCaMP6s in excitatory neurons via tetO-GCaMP6s (Jax #024742) x Camk2a-tTA (Jax #003010). We confirmed the selectivity of this approach using RNAscope (SFig. 8). ChroME (Mardinly et al., 2018) or ChroME2s (Sridharan et al., 2022) was transfected via AAV. All constructs were bicistronically linked to a nuclear localized mRuby3 used for targeting photostimulation. Excitatory specificity was ensured using either a cre dependent AAV (Syn-ChroME Addgene 170161; or CAG-ChroME2s Addgene 170163) in an excitatory specific cre line (Emx1-Cre, Jax #005628. Or SepW1-Cre MGI:5519915) or a Tta dependent AAV (Tre-ChroME or Tre-ChroME2s Addgene 170177) using the same Camk2-tTa source as above. In some cases, other cre lines (Jax #017320, or Jax #013044) were crossed to the tetO-GCaMP6s x Camk2-tTa line, with other cre dependent AAV fluorophores/indicators, those results are not a part of this study. No difference was observed between any mouse preparation (SFig. 9). Control mice had the same tetO-GCaMP6s x Camk2a-tTA without any viral injections. Mice were housed in cohorts of five or fewer in a reverse light:dark cycle of 12:12 hours, with experiments occurring during the dark phase.

### Surgery

All experiments were performed in accordance with the guidelines and regulations of the Animal Care and Use Committee of the University of California, Berkeley. For head fixation during experiments, a small custom stainless-steel headplate was surgically implanted. Briefly, adult mice (P35-P50) were anesthetized with 2-3% isoflurane and mounted in a stereotaxic apparatus. Body temperature was monitored and maintained at 37°C. The scalp was removed, the fascia retracted, and the skull lightly scored with a drill bit. Vetbond was applied to the skull surface, and the headplate was fixed to the skull with dental cement (Metabond). A fine-point marker was used to note the approximate location of bregma and the left primary visual cortex (V1; 2.7mm lateral, 0mm posterior to lambda). 2-3 burr holes were drilled using a dental drill (Foredom) with a 0.24mm drill bit (George Tiemann & Co.), and 200-300nl of AAV was injected at 50nl/min, followed by a 5+ min waiting period. A 3-3.5mm region of skull surrounding the marked V1 area was removed using the dental drill and/or a biopsy punch (Robbins Instruments). The window was replaced with three glass coverslips (two 3mm and one 5mm) and cemented into place with dental cement. Mice were given additional saline during surgery (0.3ml 0.9% NaCl). Mice received buprenorphine and meloxicam for pain management and dexamethasone to reduce brain swelling.

### Multiphoton Imaging and Stimulation Microscope

All *in vivo* experiments were performed using a setup capable of 3D scanless holographic optogenetics with temporal focusing (3D-SHOT), as described previously (Pégard et al., 2017; Mardinly et al., 2018; Bounds et al., 2022; Sridharan et al., 2022). The microscope is adapted on a Movable Objective Microscope (MOM; Sutter Instrument Co.) platform, with three combined optical paths: a 3D two-photon (2P) photostimulation path, a fast resonant-galvo raster scanning 2P imaging path, and a widefield one-photon (1p) epifluorescence/IR-transmitted imaging path, merged by a polarizing beamsplitter before the microscope tube lens and objective. Imaging was performed with a Chameleon Ultra ii (Coherent inc), and photostimulation with a Monaco40 (Coherent inc). Temporal focusing of the photostimulation beam from the femtosecond fiber laser was achieved with a blazed holographic diffraction grating (R5000626767-19311 Newport Corporation). The beam was relayed through a rotating diffuser to randomize the phase pattern and expand the temporally focused beam to cover the area of the high-refresh-rate spatial light modulator (SLM; HSP1920-1064-HSP8-HB, 1920 × 1152 pixels, Meadowlark Optics). Holographic phase masks were calculated using the Gerchberg-Saxton algorithm and displayed on the SLM to generate multiple temporally-focused spots in 2D or 3D positions of interest. The photostimulation path was then relayed into the imaging path with a polarizing beamsplitter placed immediately prior to the tube lens. As described in Mardinly et al., 2018, to limit imaging artifacts introduced by the photostimulation laser, the photostimulation laser was synchronized to the scan phase of the resonance galvos using an Arduino Mega (Arduino), gated to be only on the edges of every line scan.

### Calibration

Multiphoton activation of cells requires very precise alignment of the stimulation and the imaging system throughout a large 3D volume. Most calibration procedures assume that individual imaging and stimulation planes are parallel and flat. However, certain optical elements and subtle misalignments of the microscope can add aberrations that introduce mistargeting errors, especially at the edges of the field of view. For this reason, we improved our previous calibration approaches (Pégard et al., 2017; Mardinly et al., 2018; Oldenburg et al., 2022)with a new fully automated multiplexed 3D calibration, that accounts for arbitrary distortions in either the imaging or stimulation planes (SFig. 1A-H). We confirmed that our system is able to deliver arbitrary powers to arbitrary locations in single and multi-target holograms. As expected, we found that multitarget holograms were less efficient than single target holograms, i.e., more light is lost to diffraction. But for holograms of 3 or more targets the light intensity hitting a given target is not affected by the identity or number of other targets (SFig. 1I). For this reason, in all subsequent experiments we restrict holograms to contain at least 3 target cells.

### Holographic stimulation

Cells were targeted for stimulation based on the nuclear localized mRuby signal bicistronically linked to the opsin. Only multitarget holograms of at least 3 targets were used. Putative opsin positive cells were analyzed online using scanImage (vidrio inc) by collecting fluorescent scores around each automatically detected red nuclei. ROIs that were not holographically activatable were not included in further experiments. Online data was only used during the experiment and was not used in analyses.

To minimize the risk of off target activation, we minimize the power used per cell by first performing a ‘power test’ on each cell. In groups of 5 cells at a time, we activated each cell with five 5 ms pulses of light at powers ranging from 12.5 to 100 mW/cell. We define the stimmable power as the power in which we could elicit a significant calcium response in a given cell. ChroME, and its derivatives, are useful in that using excess power does not easily elicit more than one spike per 5ms pulse (Mardinly et al., 2018; Bounds et al., 2022; Sridharan et al., 2022). Therefore, we multiply the stimmable power by 1.1 to 1.2 to ensure more faithful response in each stimulated cell. Throughout the experiments, multi target holograms are designed such that each cell receives a distinct power based on its stimmability and the diffraction efficiencies of each spot. We further restrict analysis to exclude cells within 15 μm on the same plane or within 30 μm one plane away (30 μm spacing), as they have a risk of receiving off target light.

To confirm our resolution, we obtained physiological point spread functions in two separate experiments from a total of 26 matched cells. After the standard ‘power test’, randomly selected sets of 10 cells distributed throughout the field of view were driven as in a standard experiment. Holograms were digitally offset radially using 3 μm steps (range −3 to 30 μm from aligned), and axially in 6 μm steps (range −6 to 60 μm). Resulting fluorescence was fit with a gaussian, aligned to the peak and the full width half max (FWHM) was obtained.

### Calcium Imaging

All recordings were performed in L2/3 imaging three 800 x 800 μm planes, spaced 30μm apart, at 5.2-6.2Hz with <75mW 920nm laser light (Coherent Chameleon) using a resonant galvo system. Images were acquired using ScanImage (Vidrio Inc.) with custom stimulation control software. During recordings, animals are on a running wheel and their run speed is recorded.

Visual stimuli were presented on a 2048 x 1536 Retina iPad LCD display (Adafruit Industries) placed 10 cm from the mouse. The monitor backlight was synchronized with the galvos such that it came on only during the turnaround time, so that light from the monitor did not contaminate 2P imaging. Visual stimuli were created and presented with custom MATLAB code and Psychophysics ToolBox. Drifting gratings (50 visual degrees, 1 Hz, .08 cycles per degree, 100% contrast) of different orientations were randomly presented for 1 second each trial and interleaved with a grey-screen (“blank”) condition. Neurons with significantly different responses to visual stimuli (p<0.05, ANOVA) were considered as visually responsive.

### Online Analysis

Tuning curves and responses to photostimulation were calculated during the experiment, using a custom online implantation of CaImAn OnACID (Giovannucci et al., 2019) to perform rigid motion correction and seeded source extraction (https://github.com/willyh101/live2p). Preferred orientation (PO) was calculated as the max mean response to oriented gratings, and orientation selectivity was calculated as (PO-OO)/(PO+OO).

### Discrete Optimizer

In some experiments it was difficult to manually identify the optimal targets to create distinct ensembles that fit certain criteria, such as close and o-tuned ensembles. To overcome this challenge, we wrote a custom discrete optimizer. This optimizer selects groups of targets to stimulate from a database of eligible cells to minimize a custom cost function. As ‘cells to include’ is a discrete operation each step of our optimizer swaps one or more cells before evaluating the cost function and continuing. For a given experimental day we optimized for 3-20 ensemble of 10 cells that (1) were maximally distinct from each other, (2) minimized the number of times individual cells were included in different holograms, (3) prioritized cells activated with low light powers, (4) prioritized visually responsive cells, (5) avoided instances where two cells in the same ensemble were within 30 μm of each other, (6) spread out cells within an ensemble, (7) fit the desired spatial rules (e.g. were spatially compact vs spread out), (8) fit the desired ensemble tuning (i.e. ensemble OSI), and (9) fit the desired mean selectivity (i.e., OSI of ensemble members was high for co-tuned ensembles or low for untuned ensembles).

### Offline Analysis

Tiff files were motion corrected, cell sources (aka pixel masks) were determined, and source fluorescence was extracted using suite2p (Pachitariu et al., 2017). Pixel masks were manually categorized as ‘cells’ or ‘not cells’ and only ‘cells’ were included for analysis. For ΔF/F calculation, each cell’s detected fluorescence was first neuropil subtracted. The average fluorescence of an anulus (not containing another cell) of up to 350 pixels was considered neuropil. A neuropil coefficient (c) was calculated for each cell as described in (Pachitariu et al., 2017) and the final fluorescence was calculated as F= F_cell_ – c * F_neuropil_. F_0_, the ‘baseline’ fluorescence, was calculated with a moving average of the 10^th^ percentile of a 1000 frame window (approx. 3 minute); this moving average corrected for very slow drift in imaging conditions. ΔF/F is (F-F_0_)/F_0_.

Not all putative cells identified via red nuclei and/or online analysis were recovered by suite2p. This ‘non-matched’ population could be caused by a variety of sources, including errors in the online initial detection algorithms, errors in suit2p’s recovery, and potentially errors in the manual ‘cell’ vs ‘not cell’ determination. If too many cells of an ensemble did not match that ensemble was excluded (see exclusions). When calculating distance to a target, or spread of an ensemble, the targeted rather than recovered sets of coordinates are used.

The minimum distance to a target was defined for each cell as the minimum distance to any attempted target, regardless of if that target ‘matched’ to a suite2p detected cell.

The spread of an ensemble was calculated as the mean pairwise distance between the center of mass of each target of an ensemble calculated in 3D. A close ensemble is defined as having a mean pairwise distance < 200 μm whereas a far apart ensemble has a mean distance > 200 μm.

Tuning curves and OSIs were recalculated offline data for subsequent analysis. Ensemble OSI is defined as the OSI of the mean tuning curve from cells used in an ensemble. Mean OSI is the arithmetic mean of the OSIs from each ensemble. Co-Tuned ensembles are defined as ensembles with an ensemble OSI > 0.7 and a mean OSI > 0.5; untuned ensembles are defined as ensembles with an ensemble OSI < 0.3 and a mean OSI < 0.5.

### Exclusion Criteria

Trials were excluded if (1) the animal ran more than 6 cm/s, (2) 50% or more of the targeted cells failed to respond when driven (to at least 0.25 Z-scored Fluorescence above baseline), or (3) registration of the field of view indicates that the brain shifted more than 4.7μm (3 pixels), indicating a miss.

Cells were excluded from a given trial if (1) they were located in an off target region (15μm radially from a targeted cell, or 30μm radially from a cell one plane away), (2) they had been stimulated in the immediate preceding trial, (3) they were occluded by the stimulation artifact, or (4) the cell was categorized as ‘not cell’ or not detected via the suite2p process.

Ensembles were excluded from analysis if (1) more than 33% of the targeted cells were not detected via suite2p, (2) more than 50% of attempted stimulation trials failed (note only successful trails are included), or had fewer than 10 repetitions for either the baseline (4) or stimulation (5) conditions.

Fields of view were excluded from analysis if (1) fewer than 5% of cells were visually responsive, (2) more than 50% of trials were occurred while the mouse was running, or (3) fewer than 250 total cells were detected by suite2p.

### Determining Opsin Negative Cells

Opsin positive and opsin negative cells were identified in SepW1-Cre x CamK2a-tTA x tetO- GCaMP6s mice injected with AAV-CAG-DIO-ChroME2s-P2A-H2B-mRuby3 as described above. In addition to the typical imaging procedures, a structural image of the field of view at 1020nm was taken at the start of an experiment to identify and quantify the brightness of the nuclear mRuby3. As window preparations and imaging conditions could vary between days the mRuby3 brightness was considered a relative measure. For each field of view the top 20% brightest red nuclei were defined as opsin positive, while the 30% dimmest were considered opsin negative. Opsin negative cells often scored low integer values fluorescent counts, with many cells receiving equal scores, thus in some recordings more than 30% of cells were included.

### Mathematical model

We consider a two-dimensional neural field model of the form

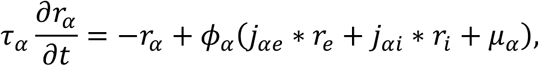

where * denotes a two-dimensional convolution in space, *α* = *e*, *i,* and *x* ∈ [0,1400] × [0,1400]μm box with periodic boundary conditions (modified from Huang et al., 2019). Since the animal is viewing a gray screen, there exists a uniform steady state for this system ***r**_ss_*. We then make use of the fact that the perturbation to the system is relatively weak, and as a result, we can approximate it as a linear perturbation around ***r**_ss_*. Linearizing the system yields

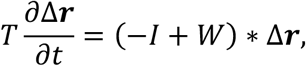

where

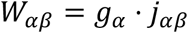

with *g_α_* being the gain set by the steady state of the system. The optogenetic stimulation is then modeled by considering

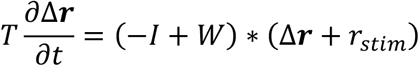

where *r_stim_* = 10 · *δ*(*x* – *x_i_*) *δ*(*y* – *y_i_*) at ten stim locations denoted by (*x_i_,y_i_*). Transforming the system into Fourier space, we can solve for Δ***r*** in steady state

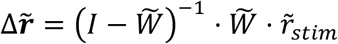

We can perform a matrix expansion of this inverse as long as the spectral radius of 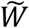 is less than one. Since V1 is in a low gain state while the animal is viewing a gray screen, the effective connection strength is very weak (i.e., *g_α_* ≪ 1), placing us within this regime. Performing this expansion yields

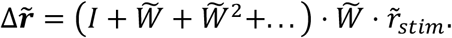

Finally, after taking the inverse Fourier transform, we find that

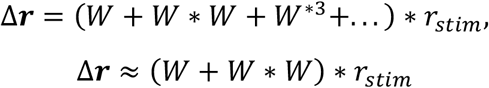

where we again made use of the weak effective connectivity to drop the higher ordered terms. Since we are only stimming and recording from excitatory neurons, we can write this approximation as a sum of monosynaptic and disynaptic excitatory terms, and a disynaptic inhibitory pathway

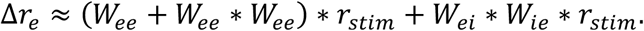

While we consider connectivity rules that depend on both space and feature, we assume that these components are independent. This allows us to write the strength of connection between neurons at coordinates (*x*_1_,*y*_1_) and (*x*_2_,*y*_2_) with feature preference *θ*_1_ and *θ*_2_, respectively, as

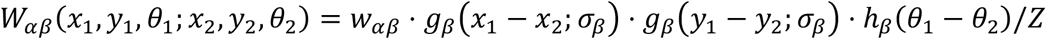

where *Z* is a normalization factor to ensure that the integral of *W_αβ_* equals *w_αβ_* The spatial dependence is given by the following sum of wrapped Gaussian distributions

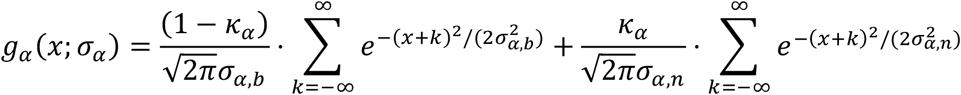

where *σ_α,b_* refers to the broad spatial component, *σ_α,n_* is the narrow spatial component, and *κ_α_* is the relative weight of each of them. The broad spatial components for both outgoing excitatory and inhibitory connections are fit according to data from (Rossi et al., 2020) (SFig. 5A-D). The parameters of the narrow component, *σ_α,n_* and *κ_α_*, are adjusted from Yu et al. (2020), chosen to capture the nearby excitation observed in Fig. 2A and SFig. 5E. Further, the boundary of the experimentally observed data regime box used in Fig. 2C was found by fitting experimental data for different bin widths (5 μm-20 μm) to the function

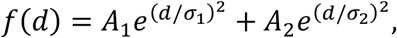

solving for the zero crossing and max activation/max suppression, and then taking the boundary to be the smallest rectangle that includes the values for all bin widths.

The models without feature-based connectivity take *h_α_*(*θ*). Otherwise, it takes the form

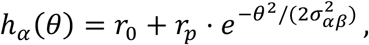

where *θ* ∈ [0,90]. The parameters for feature-based excitatory connections are also fitted according to data from (Rossi et al., 2020), whereas the inhibitory connections are adjusted to best match the observed like-to-like suppression seen in Fig. 4. After using the available data, the free parameters are the effective strength of excitatory connections, *w_ee_*, the effective strength of the inhibitory pathway, *w_eie_* = *w_ei_ · w_ie_*, the narrow spatial components parameters, and the feature base rules of the inhibitory connections.

### RNA in situ hybridization

Brain was harvested from a 6 months old tetO-GCaMP6s+/+ CamK2a-tTA+/- female, embedded in optimal cutting temperature compound (OCT, Tissue-Tek), and frozen on dry ice within 5 minutes of tissue harvest. Tissue blocks were cut into 10 μm sections using a cryostat. RNAscope ™ was performed on the sections according to the manufacturer’s instructions (RNAscope ™ Fluorescent Multiplex Kit, Advanced Cell Diagnostics). Probes used were Mm- GCaMP6s-O1, Mm-Slc32a1-C2, and Mm-Slc17a7-C3. RTU DAPI was used to stain cell nuclei and slides were mounted using Vectashield mounting medium (Vector Labs). Images were collected using LSM 880 NLO AxioExaminer confocal microscope (Zeiss) and processed using ZEN lite (Zeiss). For analysis, 300 cells with positive DAPI stating were counted in cortical Layer 2/3 and positive/negative staining of each probe was recorded for each cell. Cell with less than 10 dots per probe was presumed negative for the respective RNA.

## Data/Model Availability

The data and code will be made available upon acceptance for publication.

## Acknowledgements

This work was supported by NIH grants U19NS107613, R01EY023756, RF1MH120680, UF1NS107574 and a New York Stem Cell Robertson Fellowship and Chan Zuckerberg Biohub Investigator Award to H.A; NIH grants U19NS107613 and R01EB026953, and a grant from the Simons foundation collaboration on the global brain to B.D.; NSF Fellowship DGE 1752814 to H.A.B; Swartz Foundation Fellowship for Theory in Neuroscience, and the Burroughs Wellcome Fund’s Career Award at the Scientific Interface to G.H.; NIH Grant F31EY031977 to W.D.H.; and NIH grant K99EY029758 and the Simons Foundation collaboration on the global brain fellowship to I.A.O.

---

**SFig 1:**
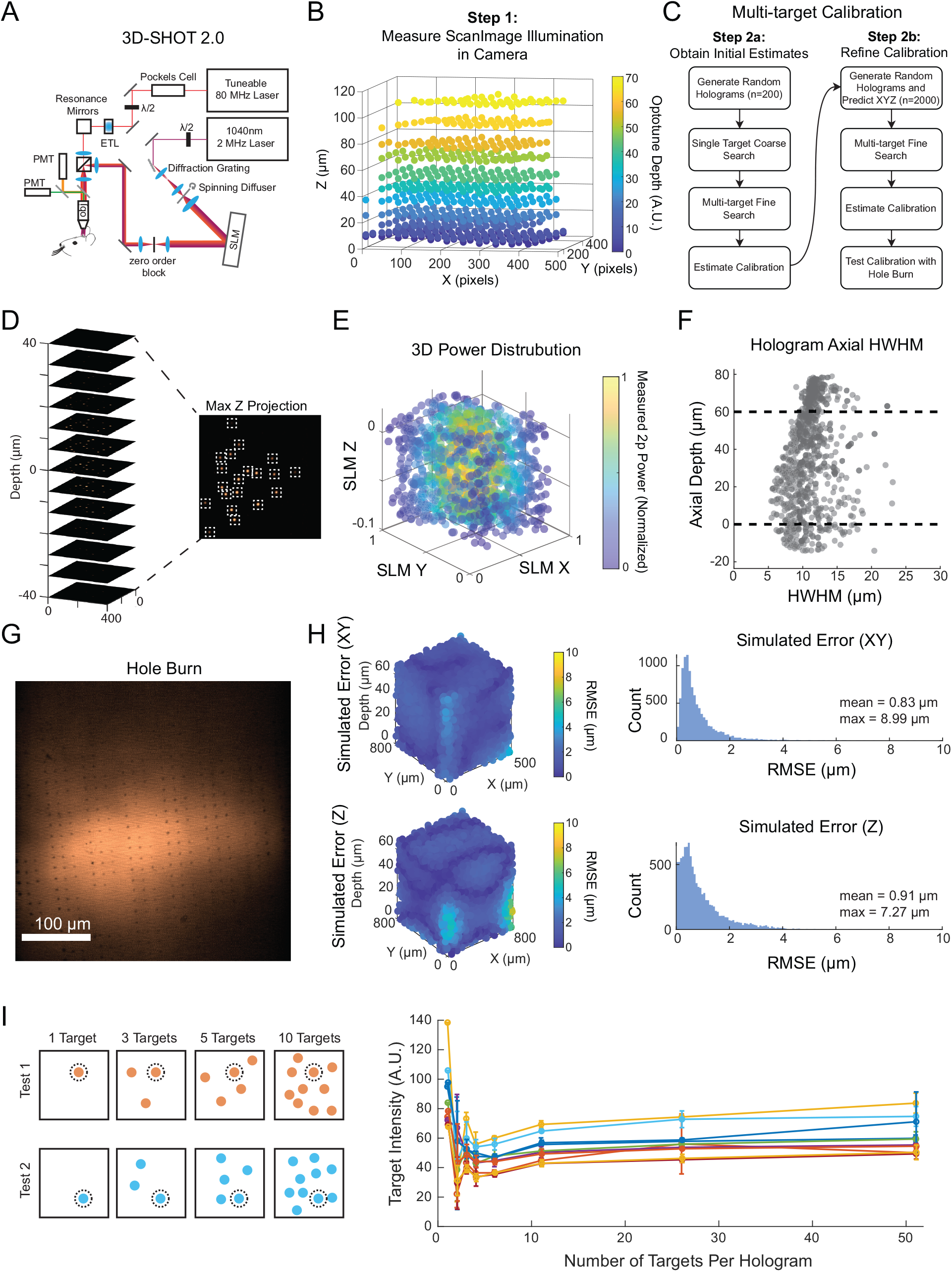
3D-SHOT2.0 and 3D spatial calibration. **A:** Microscope setup implementing simultaneous holographic optogenetics (via 3D-SHOT 2.0) and 2-photon calcium imaging. Imaging and stimulation paths are co-aligned and calibrated using the procedure outlined in this figure. **B:** First, a substage camera is used to image the ‘imaging’ fields of view in three dimensions. A thin fluorescent slide is imaged by the camera, and the microscope focus moved above and below the slide to gain depth information. The field curvature, introduced by the ETL, among other elements, was measured as a function of XY position and depth and then fit with a polynomial. Multiple ETL offsets representing the different imaging planes that might be used in an experiment are imaged separately. **C:** Second, we devised a multiplexed approach to register the position and diffraction efficiency of hologram targets (aka individual illuminated spots) throughout the Imaging FOV. To begin, the approximate XYZ position of 200 randomly targeted single-target holograms were determined by imaging a with the substage camera. Each hologram was projected onto the thin film slide, and then a z-stack of images made by defocusing the microscope. The XY position of each hologram was determined by the position on the camera, while the Z position inferred from the stack. Next, multi-target holograms comprised of the holograms imaged in the initial phase were imaged at a higher resolution to provide a more accurate XYZ position. By imaging holograms as multi-targets we were able to parallelize data acquisition and dramatically improve calibration times. Next, we use the initial 200 imaged holograms to estimate the calibration and registration of holograms in the imaging FOV. Using these initial model fits, we then generate 100 20-target holograms (for a total of 2000 data points), predict their location in the substage camera, and extract the resulting XYZ position, relative power, and HWHM throughout the entire imaging FOV. Data were fit with a polynomial to generate a general transform for any SLM and imaging coordinate. Finally, we test the spatial calibration with an automated “hole-burn”. **D.** An example of a multi-target hologram being imaged in a stack. **E:** Example power distributions in 3D (presented in arbitrary SLM units). Power distributions vary significantly throughout 3D. By modeling hologram diffraction efficiency in 3D, we can dynamically compensate power during an experiment to accurately use a greater range of the SLM. Each point represents a single imaged hologram; color represents the relative power from that hologram. **F:** Axial hologram HWHM varies as a function of depth. We co-aligned the axial position of the imaging and holographic stimulation pathways such that the holograms with smallest HWHM were positioned within the typical Imaging axial range used in experiments (dashed lines). Each point represents a single imaged hologram. **G:** Example image from the automated “hole-burn”. A unique pattern is bleached/burned onto a thin fluorescent slide, imaged in ScanImage, and the XY position of hole-burn locations is detected. This process is repeated for multiple z-planes. **H:** With the calibration in place, we simulated XY (top) and Z (bottom) error in targeting throughout the typical imaging field of view (n=10,000 simulations). Root-mean-square error (RMSE, in μm) is presented in 3D (left) and overall error distributions (right). **I:** To understand the variability in delivered power across different types of holograms, we measured the fluorescence evoked by a series of 10 test targets (dotted circle, left). Using the substage camera, we measured the intensity from the test target alone in a single-target hologram, or in a hologram that also contained 1 to 50 randomly chosen ‘distractor’ targets, 10 repeats with new distractor targets were performed per test target and hologram size. *Right,* the intensity of each test target as a function of the total number of targets in the hologram. Each color denotes a different test target, while the variability comes from different ‘distractor’ targets (presented as S.E.M).

**SFig 2:**
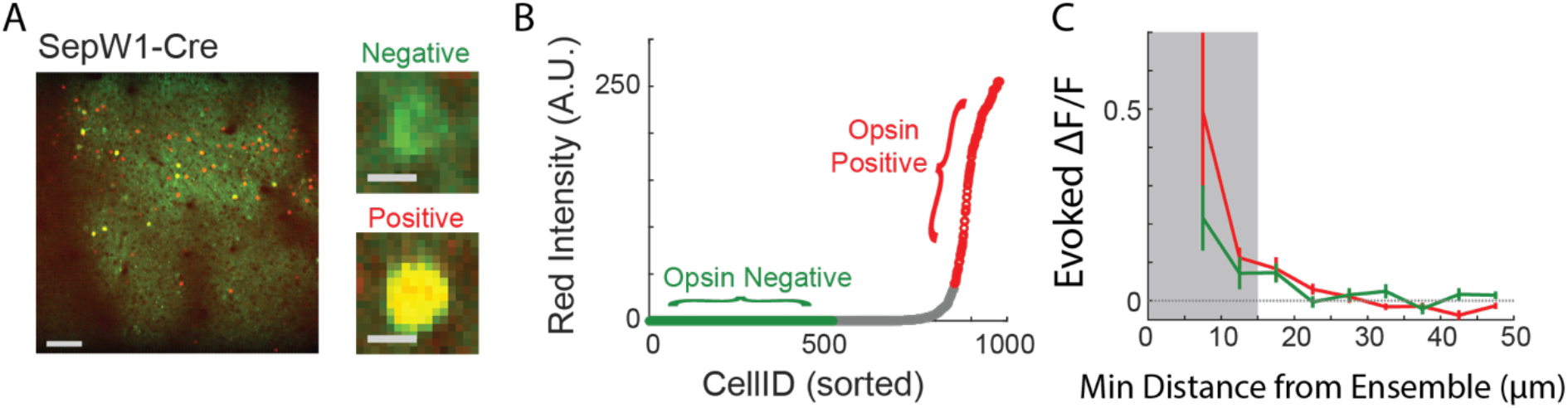
Sparsely Expressing Mice for analysis of opsin negative cells. **A:** Representative image from a FOV of a SepW1-Cre mouse with AAV CAG-DIO-ChroME2. Scale Bar 100 μm. Inset, example images of opsin negative (above) and opsin positive (below) cell. Scale Bar (10 μm). Green is from GCaMP, red from nuclear localized mRuby3 part of the opsin construct. **B:** The intensity of the red fluorophore detected for each cell in a field of view. Top 80^th^ percentile categorized as opsin positive, bottom 30^th^ percentile opsin negative (Note: red counts are integers and more than 30% of cells may have the same or lower 30^th^ percentile score). **C:** Evoked ΔF/F responses in non-targeted cells to 10 cell ensemble stimulation as a function of distance from the closest target, separated by opsin positive and opsin negative cells. Exclusion zone (15 μm) marked in grey. N= 57 ensembles, 6 FOVs, 4 Mice.

**SFig 3:**
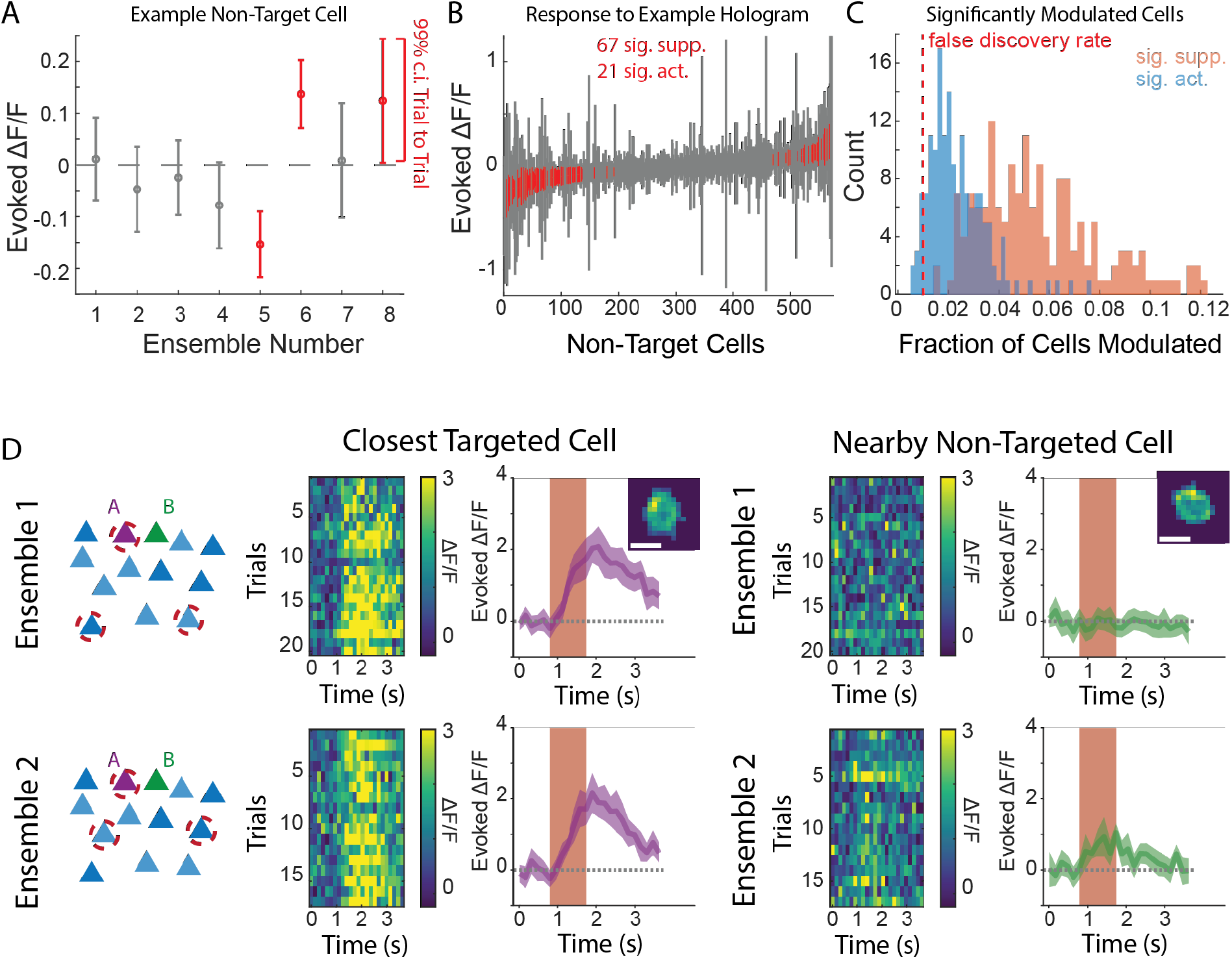
Heterogeneity in Responses to Holographic Stimulation. **A:** Evoked ΔF/F responses in a single example non-target cell to 8 distinct 10 cell ensembles stimulations from the same recording. Trial variability presented as mean ± 99% c.i.. Significantly modulated conditions (i.e., 99% c.i. excludes 0) are marked in red. **B:** All non-target cells from a given field of view response to a single example hologram. Mean ± 99% c.i. sorted by mean response. Cells that are significantly modulated (i.e., 99% c.i. excludes 0) marked in red. To this hologram 67 cells were significantly suppressed, while 21 were activated. **C:** For each stimulated ensemble, the fraction of non-target cells significantly activated (blue) or suppressed (orange) (via 99% c.i.) is presented as a histogram. False discovery rate (1%) is noted by the dotted red line. N=160 Ensembles, 18 FOVs, 13 Mice. **D:** An example pair of cells that were close to each other (<30μm) but responded differently to different, but similar, ensembles. Cell A (purple) was targeted in both Ensemble 1 (top) and Ensemble 2 (bottom). Cell B (green) was never directly targeted but was silent in ensemble 1 and driven in ensemble 2. Each row shows a schematic of the ensemble (far left), color plot of the fluorescence observed each trial for cell A (left), mean ± 95% c.i. response for cell A (middle), color plot of the fluorescence observed each trial for cell B (right), and mean ± 95% c.i. response for cell B (far right).

**SFig 4:**
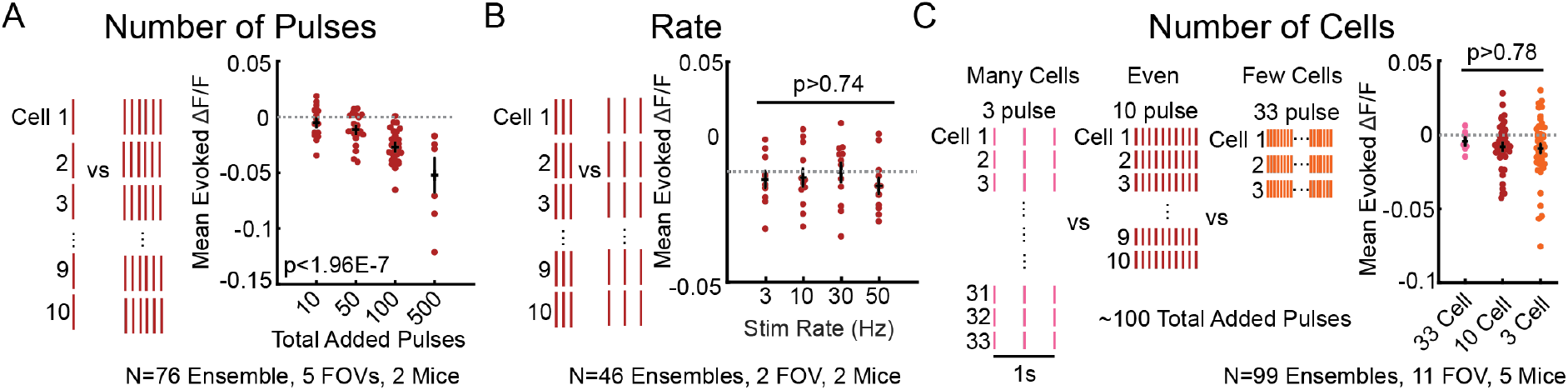
Observed decrease in fluorescence is dependent on the number of spikes added. **A:** Number of total pulses added increased overall mean suppression. Mean evoked population response to 10 cell ensemble stimulation, with 1, 5, 10 or 50 pulses per cell. P<1.97e-7 ANOVA, N=76 Ensembles, 5 FOVs, 2 Mice. **B:** Rate of stimulation did not affect the overall mean suppression. Mean evoked population response to 10 cell ensemble driven with 10 pulses at 3, 10, 30 or 50 Hz. P>0.74 ANOVA, N=46 Ensembles 2 FOVs, 2 Mice. **C:** Number of cells stimulated, when holding number of spikes constant, did not change overall suppression. Ensembles of 33 cells driven with 3 pulses, vs 10 cells driven with 10 pulses, vs 3 cells driven with 33 pulses did not recruit a differential amount of suppression. P>0.78 ANOVA, N=99 Ensembles, 11 FOVs, 5 Mice.

**SFig 5:**
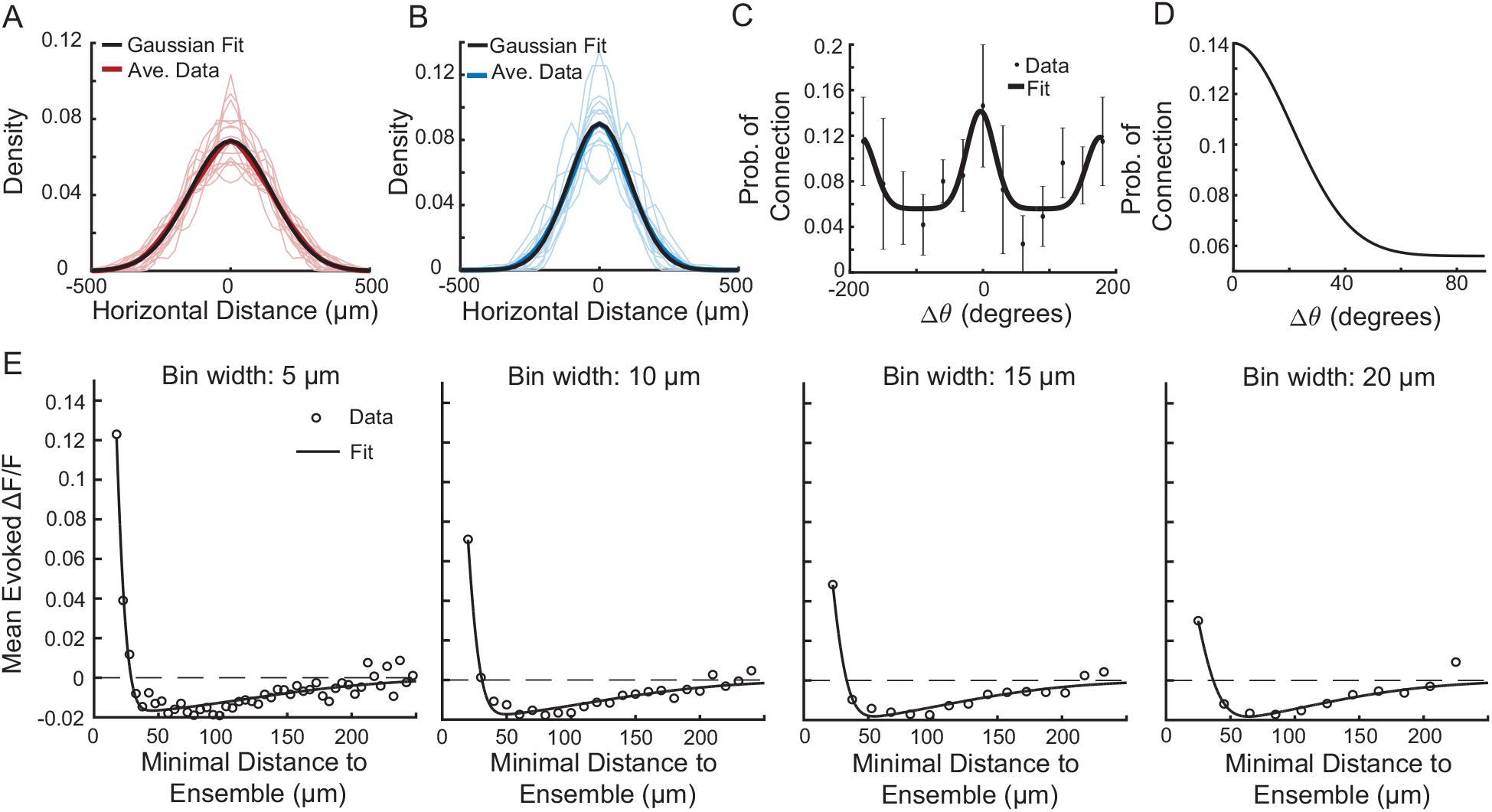
Using experimental data to fit spatial and feature spread parameters. **A:** Data from Rossi et al. (2020) used to fit the broad spatial component of E→α, for *α* = E, I (σ_*e,*b_ = 147.31 μm). **B:** Same as A, except for I→α (σ_*i,b*_ = 111.27 μm). **C:** Data from Rossi et al. (2020) to fit the feature-based connectivity rule for E→E connections (σ_ee_ = 21°, *r*_0_ = 0.056, *r_p_* = 0.084). **D:** Zoomed in portion of panel A from 0° to 90°. **E:** Non-targeted cell responses to optogenetic stimulation as a function of minimal distance to ensemble for different (open circles) fitted to a sum of Gaussian spatial functions for different bin widths. The zero-crossing, maximum, and minimum values were estimated for each fitted function and used to determine the experimentally observed data regime box used in Fig. 2C.

**SFig 6:**
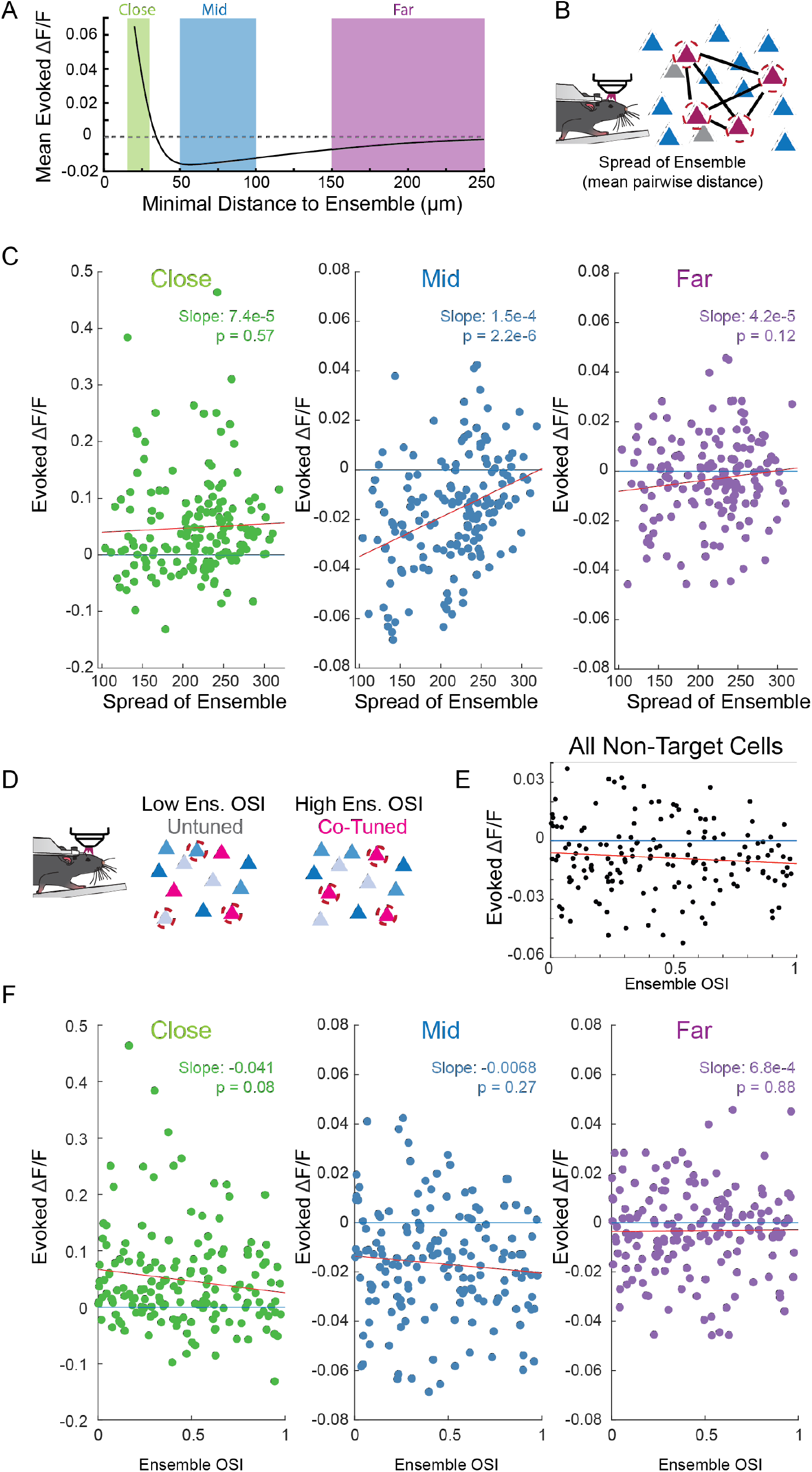
Correlation of population responses to ensembles statistics. **A:** Non-Target cells are categorized based on their proximity to a targeted cell as close (< 30 μm, green), middle (50-100 μm, blue) or far (> 150 μm, purple) cells. These categories are plotted against the predicted average response from Figure 2A. **B:** Schematic showing the mean pairwise distance of a stimulated ensemble. **C:** Population averages of evoked fluorescence from non-target cells categorized as close (green), middle (blue), or far (purple) from a targeted cell as a function of the ensemble spread. Each dot is a population response to an ensemble, N=160 Ensembles, 18 FOVs, 13 mice. Blue line 0 effect, red line linear regression fit. Slope and linear regression p value written on the plot. **D:** Schematic showing tuned vs untuned ensembles. **E:** The evoked fluorescence from all non-targeted cells regardless of distance to a stimulated cell as a function of ensemble OSI (i.e., tuning). **F:** As in C, but as a function of Ensemble OSI.

**SFig 7:**
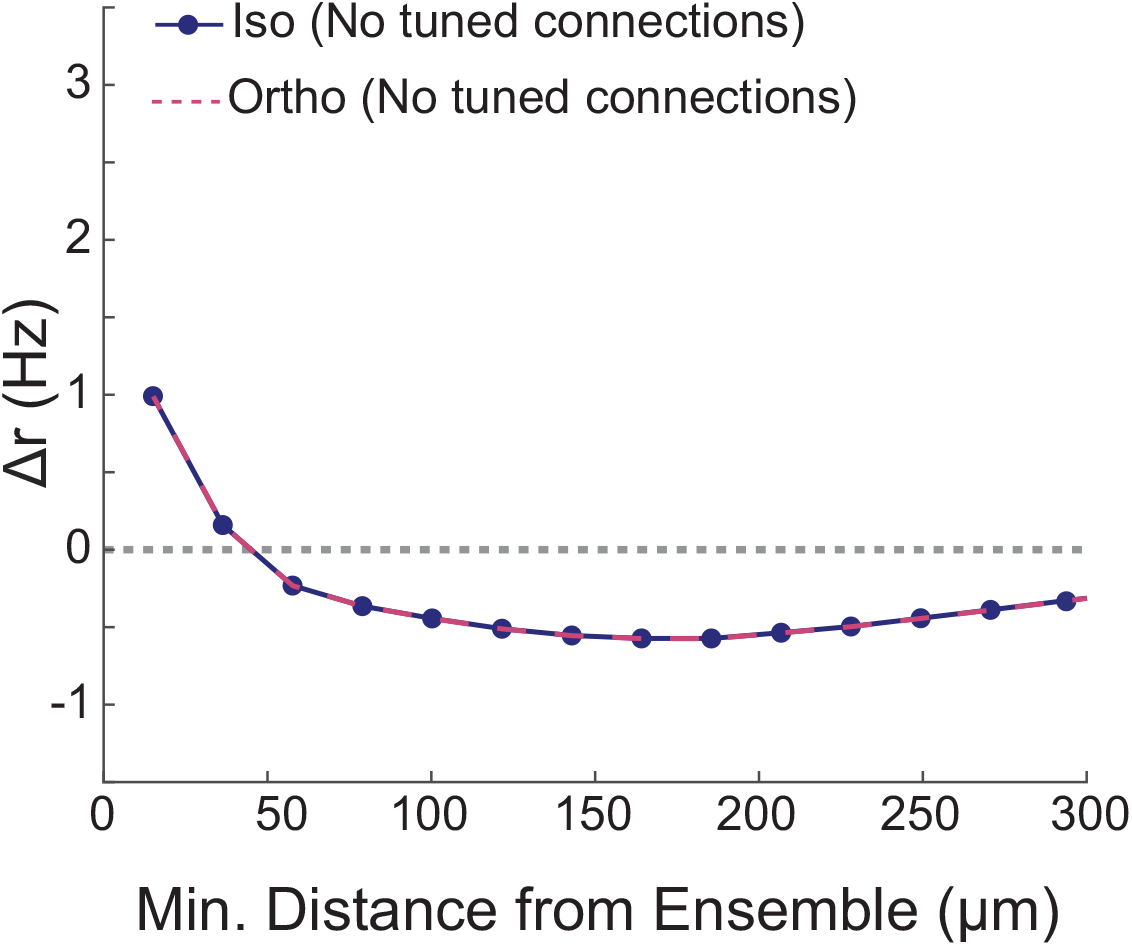
Mathematical model with no tuned connections fails to capture experimental observations. When synaptic connectivity only followed a spatial wiring rule with no specificity in orientation space (i.e., *h*_α_(*θ*)=1), there is no difference in the recruited recurrent activity of iso-oriented (solid blue) vs. orthogonally oriented (dashed red) neurons.

**SFig 8:**
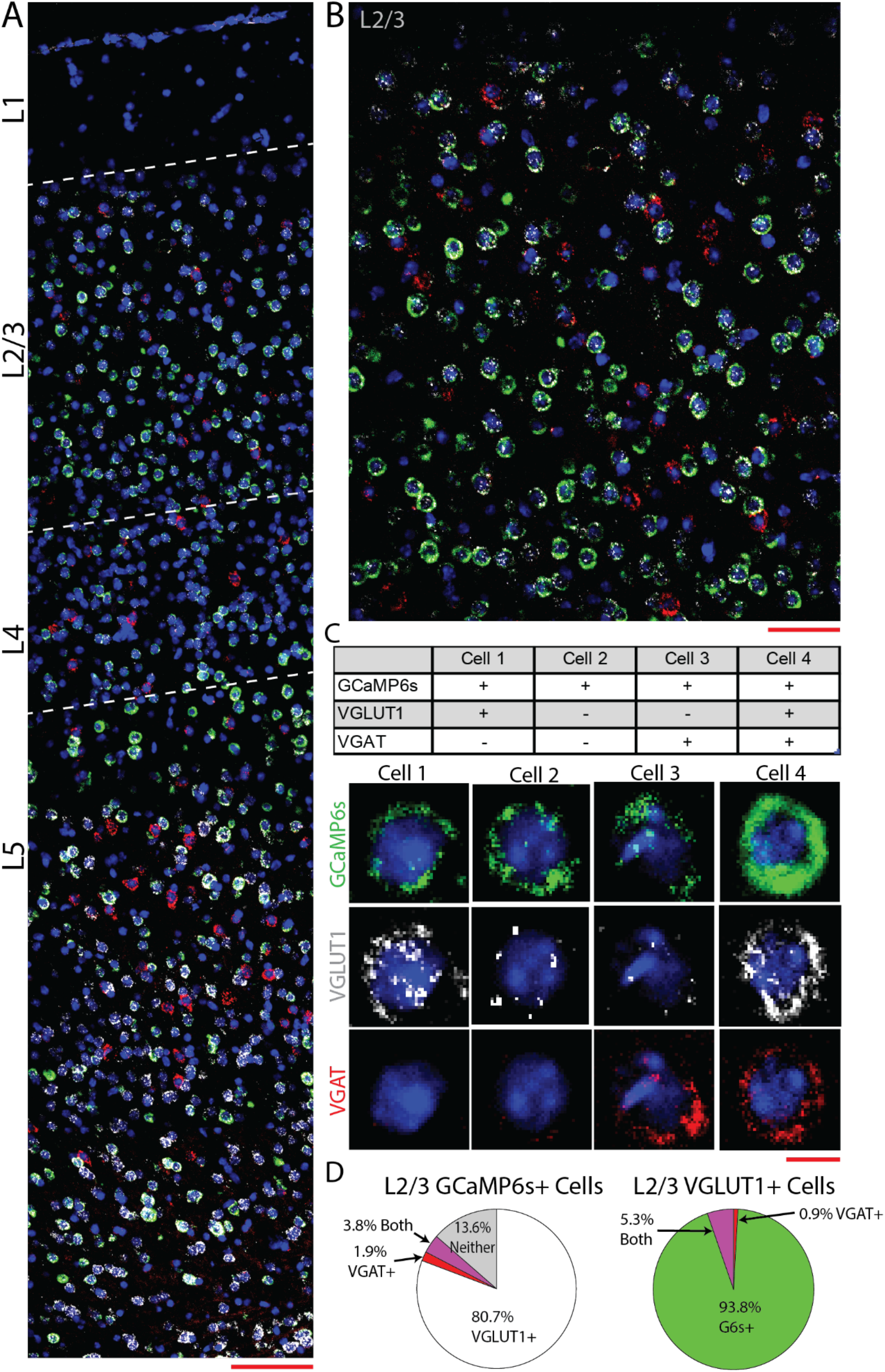
RNAscope validation of cell type specificity. **A:** A cortical section of a tetO-GCaMP6s x Camk2a-tTA mouse using RNAscope. GCaMP6s (green), VGLUT1 (white), VGAT (red), and DAPI (Blue) are labeled. Approximate division of cortical layers noted by dashed lines. Scale bar 100 μm. **B:** Enlarged view of L2/3 cortex. Scale bar 50 μm. **C:** Example L2/3 cells with expression of GCaMP6s, VGLUT1, and/or VGAT noted. Scale Bar 5 μm. **D:** Quantification of 300 DAPI positive cells, presented as either GCcaMP6s+ cells that co stained for markers of VGLUT1, VGAT, or both (left) or VGLUT1+ cells that costain for GCaMP6s, VGAT, or both (right). 84.5% of GCaMP6s+ cells stain for VGLUT1, 99.1% of VLGUT1+ cells stain for GCaMP6s.

**SFig 9:**
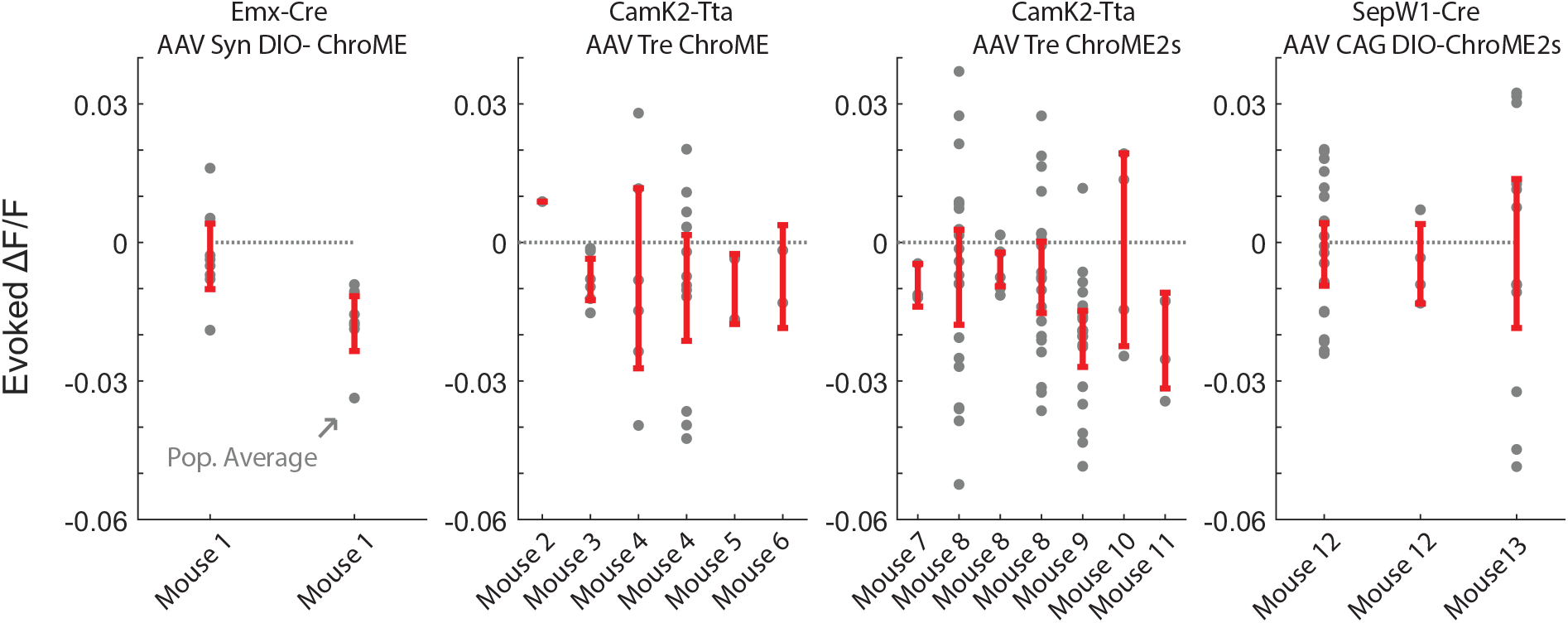
Comparison of effects by mouse and preparation. Mean population average of all non-targeted cells in response to each 10 cell ensemble stimulation (grey dot). Average ± SEM for each recording FOV in red. Divided by expression selectivity driver (Emx-cre, CamK2-tTA, or SepW1-Cre) and viral construct (Syn-DIO-ChroME, Tre-ChroME, Tre-ChroME2s, or CAG DIO-ChroME2s). Note SepW1-Cre ensembles are more likely to be spread out, due to the sparse nature.

